# Calmodulin binds to the N-terminal domain of the cardiac sodium channel Na_v_1.5

**DOI:** 10.1101/2020.06.02.129288

**Authors:** Zizun Wang, Sarah H. Vermij, Valentin Sottas, Anna Shestak, Daniela Ross-Kaschitza, Elena V. Zaklyazminskaya, Andy Hudmon, Geoffrey S. Pitt, Jean-Sébastien Rougier, Hugues Abriel

**Affiliations:** Institute of Biochemistry and Molecular Medicine, University of Bern, Bern, Switzerland; Lonza BioPharma Ltd, Visp, Switzerland; Petrovskiy Russian Scientific Center of Surgery, Moscow, Russia; Department of Medicinal Chemistry and Molecular Pharmacology, College of Pharmacy, Purdue University, West Lafayette, Indiana, 47907, USA; Cardiovascular Research Institute, Weill Cornell Medical College, New York, USA

**Author notes:** CORRESPONDENCE Prof. Dr. Hugues Abriel, MD PhD, University of Bern, Institute of Biochemistry and Molecular Medicine, Bühlstrasse 28, CH-3012 Bern, Switzerland, Twitter: @swissionchannel. These authors contributed equally (Zizun Wang, Sarah H. Vermij).

**Keywords:** Calmodulin, Sodium channels, SCN5A, Na_v_1.5 N-terminal domain, Brugada syndrome, Dominant-negative effect

## Abstract

The cardiac voltage-gated sodium channel Na_v_1.5 conducts the rapid inward sodium current crucial for cardiomyocyte excitability. Loss-of-function mutations in its gene *SCN5A* are linked to cardiac arrhythmias such as Brugada Syndrome (BrS). Several BrS-associated mutations in the Na_v_1.5 N-terminal domain exert a dominant-negative effect (DNE) on wild-type channel function, for which mechanisms remain poorly understood. We aim to contribute to the understanding of BrS pathophysiology by characterizing three mutations in the Na_v_1.5 N-terminal domain (NTD): Y87C–here newly identified–, R104W and R121W. In addition, we hypothesize that the calcium sensor protein calmodulin is a new NTD binding partner.

Recordings of whole-cell sodium currents in TsA-201 cells expressing WT and variant Na_v_1.5 showed that Y87C and R104W but not R121W exert a DNE on WT channels. Biotinylation assays revealed reduction in fully glycosylated Na_v_1.5 at the cell surface and in whole-cell lysates. Localization of Na_v_1.5 WT channel with the ER however did not change in the presence of variants, shown by transfected and stained rat neonatal cardiomyocytes. We next demonstrated that calmodulin binds Na_v_1.5 N-terminus using *in silico* modeling, SPOTS, pull-down and proximity ligation assays. This binding is impaired in the R121W variant and in a Na_v_1.5 construct missing residues 80-105, a predicted calmodulin binding site.

In conclusion, we present the first evidence that calmodulin binds to the Na_v_1.5 NTD, which seems to be a determinant for the DNE.

## INTRODUCTION

Brugada syndrome (BrS) is a genetic cardiac arrhythmia, affecting 1 in 2000 people worldwide_1_, mostly men with structurally normal hearts^2,3^. BrS is characterized by an ST-segment elevation in the right precordial ECG leads without evidence of ischemia, and patients are at increased risk of ventricular fibrillation and sudden cardiac death^2^. Approximately 20% of BrS cases are associated with mutations in *SCN5A*, the gene encoding the voltage-gated sodium channel Na_v_1.5^2^.

Na_v_1.5 is crucial for cardiac excitability as it conducts a rapid inward depolarizing sodium current (*I*_Na_) in cardiomyocytes, shaping the rapid upstroke of the action potential^2^. BrS-associated *SCN5A* pathogenic variants typically confer a loss of Na_v_1.5 function by affecting channel gating or trafficking, slowing the action potential upstroke and cardiac conduction^2^. In addition, several BrS variant channels confer a dominant-negative effect (DNE) in cellular expression systems. The DNE is defined as variant channels negatively regulating WT channels^2^. As such, when variant and wild-type channels are co-expressed in cells, the peak sodium current (*I*_Na_) is less than 50% of that in wild-type conditions. The first *SCN5A* BrS variant conferring a DNE (L325R) was described by Keller *et al.*^2^. Since then, several variant channels in the Na_v_1.5 N-terminus domain (NTD) have been shown to exert a DNE on WT channels, including R104W and R121W^2,11^.

The mechanisms underlying the DNE of NTD Na_v_1.5 variants remain unknown. Channel-channel interactions seem to be crucial for the DNE phenomenon, which is likely mediated by Na_v_1.5 interacting proteins^2,12^. Only when we identify molecular mediators of the DNE at the NTD, we can explain the functional heterogeneity of NTD variants and ultimately identify therapeutic targets for BrS patients. In this study, we aim to elucidate the mechanisms underlying the DNE of the three N-terminal mutants Y87C, R104W and R121W. We hypothesize that the calcium-binding protein calmodulin (CaM) is a yet-unknown N-terminal interaction partner. CaM is well-known to regulate voltage-gated sodium (Na_v_) and calcium (Ca_v_) channel functions^2,14^. Its interaction with Na_v_ and Ca_v_ C-terminal domains (CTD) is especially well established^2,15^. In the Ca_v_1.2 NTD, two CaM binding sites have been described^2,17^, while CaM interaction with the Na_v_1.5 NTD remains unexplored.

Here, in BrS probands of a Russian family, we identify the new Na_v_1.5 NTD variant Y87C, which exerts a DNE on WT channel function. Using the TsA-201 cell expression system in patch-clamp and biochemistry experiments, we confirm R104W exerts a DNE, but R121W surprisingly does not. CaM is identified as a new Na_v_1.5 N-terminal binding partner. CaM binds WT, Y87C, and R104W NTDs, but only weakly to R121W NTD and NTD lacking amino acids 80-105. As such, the ability of CaM to bind the Na_v_1.5 NTD correlates with the occurrence of the DNE. Lastly, we show a reduction in fully glycosylated bands of all three Na_v_1.5 variants compared to WT both at the surface and in whole-cell lysates.

## METHODS

### GENETIC ANALYSES

We obtained the informed consent from probands and their family members for genetic investigations in accordance with the Helsinki declaration. For the *SCN5A* gene, coding and adjacent intronic areas were sequenced using PCR-based bi-directional Sanger sequencing. The prevalence of the rare genetic variant c.206A>G(Y87C) was determined using allele-specific PCR in an ethnically matched group of 150 healthy volunteers, and in the public database gnomAD (https://gnomad.broadinstitute.org/). Pathogenicity assessment was performed in accordance with ACMG guidelines (2015)^2^. The potential effect of the variant was tested *in silico* using PolyPhen2.0 (http://genetics.bwh.harvard.edu/pph2/), Provean (http://provean.jcvi.org/), and MutationTaster (http://www.mutationtaster.org/) tools.

### cDNA CONSTRUCTS, CELL CULTURE, AND TRANSFECTIONS

In the cDNA templates pcDNA3.1-*SCN5A*-WT, pcDNA3.1-S-tag-*SCN5A*-WT-NTD (a kind gift from Dr. Nathalie Neyroud, INSERM, Paris, France), and pcDNA3.1-3X-FLAG-*SCN5A*-WT, the following Na_v_1.5 variants were introduced (GenScript, NJ, USA): Y87C, R104W, R121W, and Δ26, in which amino acid residues 80-105 were deleted. S-tag-*SCN5A*-WT-NTD encodes the 131-amino-acid-long Na_v_1.5 NTD. The Ca_v_1.2-WT-NTD and the homologous Ca_v_1.2-R144W-NTD constructs were generated by replacing the *SCN5A* sequence in the pcDNA3.1-S-tag-*SCN5A*-WT-NTD cDNA with the Ca_v_1.2 NTD sequence. Na_v_1.5-encoding cDNA corresponded to the transcript NM_000335.5 (human; Genbank) with the amino acid variant T559A, for which no functional consequences have been reported. Ca_v_1.2-encoding cDNA corresponded to X15539.1 (cardiac isoform, rabbit; Genbank). Primers designed for mutagenesis are available upon request. All cDNAs were validated by sequencing.

We obtained human embryonal kidney (TsA-201) and monkey kidney cells (COS) from the American Type Culture Collection (ATCC, VA, USA). Cells were cultured at 37°C with 5% CO^2^ in Dulbecco’s modified Eagle’s culture medium (Gibco, Thermo Fisher Scientific, MA, USA), which we supplemented with 2 mM glutamine and 10% heat-inactivated fetal bovine serum. Cells were kept in culture up to 20 passages for experimental use. All transfections were performed with JET PEI (Polyplus Transfection, Illkirch, France) following the manufacturer’s instructions.

For electrophysiological recordings, we cultured TsA-201 cells in 35 mm petri dishes. We transfected with 128 ng *SCN5A* cDNA to mimic a non-saturated homozygous state. To mimic the heterozygous state, cells were co-transfected with 64 ng pcDNA3.1-*SCN5A*-WT and 64 ng of a pcDNA3.1-*SCN5A* variant (Y87C, R104W, R121W, or Δ26) or empty vector (pCDNA3.1-Zeo(+)). All transfections additionally contained 64 ng pIRES-hβ1-CD8 cDNA, encoding the human sodium channel β^2^-subunit and the CD8 receptor.

For biochemical experiments, we cultured TsA-201 cells in 100 mm petri dishes. To assay cell surface biotinylation, we transfected cells with 768 ng pcDNA3.1-*SCN5A*-WT or one of the selected variants, and 364 ng pIRES-hβ1-CD8. For calmodulin pull-down experiments, we transfected cells with 2 μg pcDNA3.1-S-tag-*SCN5A*-WT-NTD or one of the selected variants.

For proximity ligation assays (PLAs), we cultured COS cells in µ-Slide 8-Well Grid-500 ibiTreat (ibidi, Gräfelfing, Germany) and transfected them with 120 ng pcDNA3.1-S-tag-*SCN5A*-WT-NTD, one of the variants (Y87C, R104W, R121W, or Δ26), or empty vector.

### RAT NEONATAL CARDIOMYOCYTE ISOLATION, CULTURE, AND TRANSFECTION

All institutional experimental guidelines for animal handling were met. The Veterinary Office of the Canton of Bern, Switzerland, approved our experiments. We isolated primary rat neonatal cardiomyocytes (RNCs) and cultured them at 37°C with 1% CO^2^.^2^ RNCs were seeded on laminin-coated (Sigma, MO, USA) µ-Slide 8 Well Grid-500 ibiTreat dishes (ibidi). Cells were transfected 24 h after cell seeding using Lipofectamine 3000 (Invitrogen, CA, USA) To study the subcellular localization of Na_v_1.5 full-length channels in the homozygous state, we transfected RNCs with 400 ng pcDNA3.1-3X-FLAG-*SCN5A*-WT or one of the variants Y87C, R104W, or R121W. To study their subcellular localization in the heterozygous state, we transfected RNCs with 200 ng pcDNA3.1-GFP-*SCN5A*-WT and 200 ng of one of the variants. All conditions additionally contained 100 ng pDsRed2-ER (Takara Clontech, Kusatsu, Japan) as an endoplasmic reticulum (ER) marker.

### CALMODULIN BINDING PREDICTION

To computationally predict CaM binding to Na_v_1.5 NTD, we used the CaM target database (http://calcium.uhnres.utoronto.ca/ctdb/ctdb/sequence.html). Scores are normalized on a scale from 0 to 9, 0 being low and 9 being high probability of CaM binding.

### BIOCHEMICAL EXPERIMENTS

#### Peptide SPOTS array

Peptides were synthesized using standard 9-fluorenylmethoxy carbonyl (Fmoc) protected and activated amino acids (Anaspec, CA, USA) on prederivatized cellulose membranes (Intavis AG, Cologne, Germany) using an Intavis robot.^2^ Human Na_v_1.5 (Q14524 (SCN5A_Human) isoform 1) N-terminal domain 1-131 was tiled as 15 amino acid peptides with a 13-amino acid overlap (two amino acids skipped between consecutive peptides). The membrane was briefly wet with dimethylformamide (DMF) and peptides were labeled with bromophenol blue (1%) for annotation of the array pattern at the completion of synthesis as described previously.^2,21^ The peptides on the array were then de-protected with two washes of 88% trifluoroacetic acid, 2% triisopropylsilane, 5% phenol and 5% water, for 90 min each at room temperature. Each de-protection step was followed by three washes with dichloromethane, three washes with DMF, and three washes with ethanol and the membranes dried. Before binding experiments, the membrane was wet by exposure to ethanol for 15 min, followed by DMF for 15 min and three 15 min washes with phosphate-buffered saline (PBS), all while shaking at room temperature. The hydrated membrane was then equilibrated for 30 min in 20 mM Tris-HCl pH 7.4, 200 mM NaCl, 1 mM EDTA, and 0.1% Tween-20, before the membrane was blocked in the same buffer plus 5% bovine serum album (BSA) for 30 min. The binding buffer used in the CaM binding reaction was identical to membrane equilibration buffer, except 5 mM EGTA was included in the buffer along with 100 nM of DyLight680-labelled CaM; mutant Cys-75 CaM purified as described previously^2,23^ and labeled with a sulfhydryl reactive maleimide fluorophore as described by the manufacturer (Fisher #46618). After 5 min at room temperature, the apoCaM binding reaction was washed in binding buffer (3×15 min including plus 5 mM EGTA) before the membrane was imaged using a Li-Cor Imaging Station (Li-Cor Biosciences, Bad Homberg, Germany).

#### Cell surface biotinylation assay

We used a cell surface biotinylation assay to study protein expression at the plasma membrane. 48 h after transfection, TsA-201 cells transiently expressing Na_v_1.5 constructs were supplied with 0.5 mg/mL EZ-link™ Sulfo-NHS-SS-Biotin (Thermo Fisher Scientific) in PBS for 15 min at 4°C. Then we washed the cells twice with 200 mM glycine in cold PBS to inactivate and remove excess biotin, respectively. Cells were taken up in lysis buffer (composed of [mM] HEPES 50 pH 7.4, NaCl 150, MgCl_2_ 1.5, EGTA 1 pH 8; 10% glycerol, 1% Triton X-100 (Tx100), Complete protease inhibitor [Roche, Basel, Switzerland]) for 1 h at 4°C. Cell lysates were centrifuged at 16,100 rcf at 4°C for 15 min. We determined protein concentration of the supernatant with Bradford assay (Bio-Rad)^2^, and incubated lysate equivalent to 2 mg of protein with 50 μL Streptavidin Sepharose High-Performance beads (GE Healthcare, IL, USA) for 2 h at 4°C, then washed the beads three times with lysis buffer, and eluted them with 30 μL of 2X NuPAGE sample buffer plus 100 mM DTT (37°C for 30 min). Input fractions were resuspended in NuPAGE Sample Buffer (Invitrogen) plus 100 mM dithiothreitol (DTT) and incubated at 37°C for 30 min.

#### Calmodulin pull-down assay

We used the calmodulin pull-down assay to determine whether Na_v_1.5 and Ca_v_1.2 constructs bind calmodulin. 48 h after transfection, TsA-201 cells were lysed in lysis buffer with 100 nM free Ca^2+^ for 1 hour at 4°C. The free Ca^2+^ concentration was calculated based on an established method^2^. Lysed samples were centrifuged at 16,100 rcf at 4°C for 15 min. We quantified supernatant protein concentration with the Bradford assay. After washing calmodulin and control Sepharose 4B beads (GE Healthcare) twice with lysis buffer, we mixed lysate corresponding to 60, 180, or 540 μg protein with the beads. Beads and lysate mixture incubated at 4°C on a wheel for 3 hours. We washed the beads three times with lysis buffer and eluted proteins from the beads in 30 µL 2X NuPAGE sample buffer with 100 mM DTT at 55°C (15 min).

#### SDS-Page and western blot

Protein samples were loaded on pre-casted 4-12% Bis-Tris acrylamide gradient gels (Invitrogen), run at 60 V for 30 min and at 200 V for 45-120 min, and transferred to nitrocellulose membranes (Bio-Rad, CA, USA) with the TurboBlot dry blot system (Bio-Rad). Western blots were performed with the SNAP i.d. system (Millipore, Zug, Switzerland). Briefly, membranes were blocked with 1% BSA, incubated with primary antibodies diluted in PBS + 0.1% Tween20 for 10 minutes, washed four times with PBS + 0.1% Tween20, incubated with secondary antibodies diluted in PBS + 0.1% Tween20, washed again four times with PBS + 0.1% Tween20 and three times with PBS. Fluorescent signals were detected on an Odyssey infrared imaging system (Li-Cor). We determined protein content by quantifying band fluorescence densities with the Image Studio Lite software version 5.2 (Li-Cor). The following primary antibodies detected the proteins of interest: mouse anti-α-CaMKII (1:3000; BD, NJ, USA; 611292); rabbit anti-α-actin (1:3000, Sigma, A2066); mouse anti-S-tag (1:1500, Sigma, SAB2702204); rabbit anti-Na_v_1.5 (1:150, generated by Pineda, Berlin, Germany); and mouse anti-alpha-1 sodium-potassium ATPase (1:3000; Abcam, Cambridge, UK; ab7671). Secondary IRDye_®_ antibodies (Li-Cor) were 680RD goat anti-mouse (1:15000; 926-68070), 800CW goat anti-mouse (1:15000; 926-32210), and 800CW goat anti-rabbit (1:15000, 925-32211).

### QUANTITATIVE REAL-TIME PCR (RT-QPCR)

We isolated total RNA from TsA-201 cells transiently expressing pcDNA3.1-hSCN5A-WT, Y87C, R104W, R121W, or Δ26 with TRIzol reagent (Invitrogen) following the manufacturer’s instructions. We used 2 µg RNA for reverse transcription with the High-Capacity cDNA Reverse Transcription Kit (Applied Biosystems, CA, USA). The cDNA was diluted (1:10) in H^2^O for qPCR with the TaqMan Gene Expression Assay (Applied Biosystems). qPCR conditions in the ViiA 7 Real-Time PCR System (Thermo Fisher Scientific) were as follows: activation 2 min at 50°C; hold 20 s at 95°C; 40 cycles of denaturation 3 s at 95°C and annealing 60 s at 60°C. We normalized the expression levels of human *SCN5A* (HS00165693_m1) to the reference gene *GAPDH* (HS99999905_m1), and then to the *SCN5A* WT expression using the 2_-ΔΔCt_ method.^2^

### ELECTROPHYSIOLOGICAL RECORDINGS

Whole-cell currents were recorded 48 hours after transfection using the patch-clamp technique with an Axopatch 200B amplifier (Molecular Devices, Wokingham, United Kingdom) at room temperature (25 ± 1°C).^2^ The recorded *I*_Na_ was filtered at 5 kHz by a low-pass filter (HumBug, Quest Scientific, BC, Canada) at a sampling rate of 20 kHz per signal. Leak current subtraction was applied with a P/4 protocol. We used the DMZ-universal puller (program 10, Zeitz, Martinsried, Germany) to pull the patch pipettes (World Precision Instruments, Friedberg, Germany) to a resistance of 2.0 to 4.5 MΩ. The internal pipette solution contained (mM) CsCl 60, aspartic acid 50, CaCl_2_ 1, MgCl_2_ 1, HEPES 10, EGTA 11, Na_2_ATP 5. pH was adjusted to 7.2 with CsOH. The external solution contained (mM) NaCl 25, NMDG-Cl 105, CsCl 5, CaCl_2_ 2, MgCl_2_ 1.2, HEPES 10, glucose 20. pH was adjusted to 7.4 with HCl.

We used the voltage clamp mode to obtain current-voltage relationships (I-V curves). Sodium current density (pA/pF) was calculated by dividing peak current by cell capacitance using the Axon™ pCLAMP™ 10 Electrophysiology Data Acquisition & Analysis Software, Version 10.7.0.3 (Axon Instruments, CA, USA). I-V curves were fitted with the equation *y* = *g*(*V*_m_ - V_rev_) / ((1 + exp[(*V*_m_ - *V*_2/2_) / K])). Activation and steady-state inactivation (SSI) curves were fitted with the Boltzmann equation *y* = 1 / (1 + exp[(*V*_m_ - *V*_2/2_) / K]), where *y* is the normalized current or conductance at a given holding potential, *V*_m_ is the membrane potential, *V*_rev_ is the reversal potential, *V*_2/2_ is the potential at which half of the channels are activated, and K is the slope factor.

### IMMUNOFLUORESCENCE AND IMAGING

#### Immunocytochemistry

To investigate Na_v_1.5 subcellular localization, we performed immunocytochemistry on transiently transfected RNCs. Two days after transfection, RNCs were fixed with cold acetone at −20°C (10 min), washed three times with PBS, and permeabilized with blocking buffer (1% BSA, 0.5 % Tx100 and 10% normal goat serum in PBS) for 30 min at room temperature. Primary antibodies diluted 1:100 in incubation buffer (1% BSA, 0.5% Tx100 and 3% goat serum in PBS) were applied overnight at 4°C. We used the following primary antibodies: mouse anti-FLAG (Sigma) to detect FLAG-Na_v_1.5 and rabbit anti-GFP (Thermo Fisher, G10362) to detect GFP-Na_v_1.5. Then, cells were washed twice with PBS and incubated for 45 mins at room temperature with AlexaFluor secondary antibodies (goat anti-rabbit 405 (Thermo Fisher Scientific)) diluted 1:200 in incubation buffer. We stained the nuclei with DAPI or NucRed dead 647 ReadyProbes Reagent (2 drops/mL, Thermo Fisher Scientific). Mounting medium (EMS shield with Dabco; EMS, PA, USA) was added to preserve fluorescence signals.

#### Proximity ligation assay

To determine whether two proteins were close *in situ*, we performed PLAs using the Duolink_®_ Starter Kit Mouse/Rabbit (Sigma) following the manufacturer’s instructions. We transiently transfected COS cells with pcDNA3.1-S-tag-*SCN5A*-WT, -WT NTD and Δ26 NTD. 48 hours after transfection, we fixed them with acetone at −20°C for 10 min. The cells were washed three times with PBS before they were permeabilized with blocking buffer (1% BSA, 0.5 % Tx100 and 10% goat serum in PBS) for 30 min at room temperature. Cells were incubated with primary antibodies diluted in incubation buffer (1% BSA, 0.5% Tx100, and 3% goat serum in PBS) overnight at 4°C. We used the following primary antibodies: rabbit-anti-S-tag (for S-tagged NTD, 1:100, Abcam, ab180958) and mouse anti-calmodulin (1:100, Merck, 05-173); or mouse anti-S-tag (for S-tagged NTD, 1:1500, Sigma, SAB2702204) and rabbit anti-Na_v_1.5 (1:150, generated by Pineda, Berlin). After washing the cells 3X with PBS, we incubated them with anti-rabbit PLUS and anti-mouse MINUS PLA probes (1 hour at 37°C) diluted in Duolink_®_ incubation buffer (1:200; Sigma), followed by three washes with PBS. We followed the manufacturer’s instructions for probe ligation and signal amplification. Duolink_®_ in situ mounting medium with DAPI was added to stain cell nuclei and preserve fluorescence signals.

#### Image acquisition and post-acquisition analysis

Confocal images were obtained with an inverted laser-scanning microscope (LSM 880, ZEN 2.1, Zeiss, Oberkochen, Germany) with a Plan-Apochromat 63X/1.40 Oil DIC objective. We collected COS cell images in the confocal mode and analyzed them with the Duolink_®_ ImageTool (Sigma) following the manufacturer’s instructions. RNC images were collected in the Airyscan super-resolution mode with the pinhole set at 1.25 Airy unit, pixel size was 35×35 nm. We used ZEN 2.1 software for processing raw Airyscan data, and the IMARIS coloc tool (IMARIS 9.3.1 software, Bitplane, Zürich, Switzerland) to quantify the colocalization between signals of two channels within the predefined ROI (region of interest, defined as the cell area). We set an automatic threshold per channel. If the automatic threshold could not be defined, we set the threshold at 30%.

### DATA AND STATISTICAL ANALYSIS

Data are represented as means ± SEM and are compared to the wild-type condition unless otherwise indicated. Statistical significance was calculated with 2-tailed Student’s *t*-test if data were normally distributed or by the Mann-Whitney U-test if data were not (Prism version 7.04; GraphPad, CA, USA). We considered a *p*-value <0.05 to indicate a statistically significant difference.

## RESULTS

### CLINICAL DESCRIPTION OF THE NEW NAV1.5 VARIANT Y87C IN BRS PATIENTS

We identified the novel missense variant c.260A>G (Y87C) in exon 2 of the *SCN5A* gene in two family members of the Russian proband family and present their pedigree chart in **Supplementary Figure 1A**. We found this Y87C variant neither in 150 ethnically matched controls, nor in the public database gnomAD (https://gnomad.broadinstitute.org/). This substitution occurred at a highly conservative position of the NTD of Na_v_1.5 (**Supplementary Figure 1D**). Three independent *in silico* tools predicted that Y87C has a deleterious effect on protein function: in the absence of functional data it was qualified as a Class III (variant of unknown significance, VUS) variant.

The 49-year-old male proband (II.2) and his eldest son of 23 years (III.2) had a characteristic spontaneous Brugada pattern on their ECGs (**Supplementary Figure 1B**). His younger 13-year-old son (III.3) has not shown any syncope or Brugada pattern on his ECG (**Supplementary Figure 1C**), and did not undergo genetic testing yet. The proband’s father (I.2) was unavailable; at the last report (age 65) he reported no syncope history. His granddaughter (IV.1) as well as his mother (I.2) have a normal ECG and no complaints. The proband was asymptomatic without history of syncope or family history of sudden cardiac death. His ECG revealed a permanent spontaneous Brugada pattern, first-degree atrio-ventricular block (PR-interval up to 240 ms), and a negative T-wave in the V1 lead (**Supplementary Figure 1B**). During nine years of follow-up, the proband repeatedly complained of chest pain after alcohol consumption. Consequently, he was hospitalized several times with an initial diagnosis of acute myocardial infarction (MI), as his ECG showed an increasing elevation of the ST-segment; however, his ECGs never showed dynamic changes specific to MI, and troponin elevation or any other biochemical MI markers were never detected. His ECG went back to the usual Brugada-like shape within 1-2 days after alcohol abstinence. Proband underwent detailed clinical examination, and the only extracardiac complaint was endogenous depression for which he required medication for many years.

The eldest son of the proband had a spontaneous Brugada pattern similar to his father’s. He had no syncope history. During nine years of follow-up, he remained sober and physically active.

Neither the proband nor his eldest son took any anti-arrhythmic medication or received any intervention. Taking into account the functional results, this variant was re-classified to Class IV (Likely pathogenic).

### NA_V_1.5 Y87C AND R104W NEGATIVELY REGULATE WT CHANNEL FUNCTION, BUT R121W DOES NOT

To investigate the consequences of the Y87C variant on Na_v_1.5 function, we used the patch-clamp technique to record the sodium current *I*_Na_ of these channels. We transiently transfected TsA-201 cells with the Na_v_ β^2^-subunit and either 100% Na_v_1.5 WT or Y87C, or with 50% WT and either 50% empty vector or 50% Y87C to mimic the heterozygous state of the patients (**Figure 1**). In the homozygous state, the Y87C peak-current density was 65.9 ± 9.5% smaller than WT (**Figure 1A,B,D**), while the half-maximal activation potential (V_2/2_) of the Y87C activation curve shifted 5.9 ± 1.3% in the depolarized direction while inactivation did not shift compared with WT (**Figure 1C, Table 1**). In the heterozygous state, co-expressing WT with Y87C led to a significant *I*_Na_ peak-current decrease of 25.1 ± 11.0% when compared with WT + empty vector (**Figure 1D**). Taken together, these data show that cells expressing only Y87C channels conduct less whole-cell sodium current than those expressing WT channel, which the shift in activation curve only partly explains. Moreover, Y87C channels exert a DNE over WT channels, as WT + Y87C conduct ∼50% less *I*_Na_ than WT + empty vector (**Figure 1D**).

**Table 1.** Biophysical properties of sodium currents conducted by wild-type and Y87C Na_v_1.5 channels. Peak sodium current (*I*_Na_) densities are extracted from the respective IV curves. WT, wild type. Data are presented as mean ± SEM and are compared to the respective wild-type values. *, *p* < 0.05; **, *p* < 0.01; **, *p* < 0.005; ***, *p* < 0.001; ****, *p* < 0.0005. Numbers of cells are indicated in parentheses.

| | $I_{Na}$ density | Steady-state inactivation | | Activation | |
| --- | --- | --- | --- | --- | --- |
| | pA/pF | k | $V_{1/2}$ (mV) | k | $V_{1/2}$ (mV) |
| Nav1.5 WT | -94.8±9.3 (n=8) | 5.1±0.1 (n=17) | -76.4±0.7 (n=17) | 7.7±0.3 (n=10) | -22.5±1.0 (n=10) |
| Y87C | -28.9±4.5 (n=11)**** | 5.1±0.2 (n=12) | -75.2±0.6 (n=12) | 9.4±0.3 (n=11)*** | -16.6±0.8 (n=11)*** |
| WT + empty v. | -51.1±13.5 (n=9)** | 5.0±0.1 (n=14) | -75.6±0.6 (n=14) | 8.2±0.2 (n=10) | -19.6±1.1 (n=10) |
| WT + Y87C | -26.0±4.3 (n=8)**** | 5.2±0.2 (n=10) | -75.7±0.8 (n=10) | 8.9±0.3 (n=8)** | -19.3±0.6 (n=8)* |

**Figure 1.**
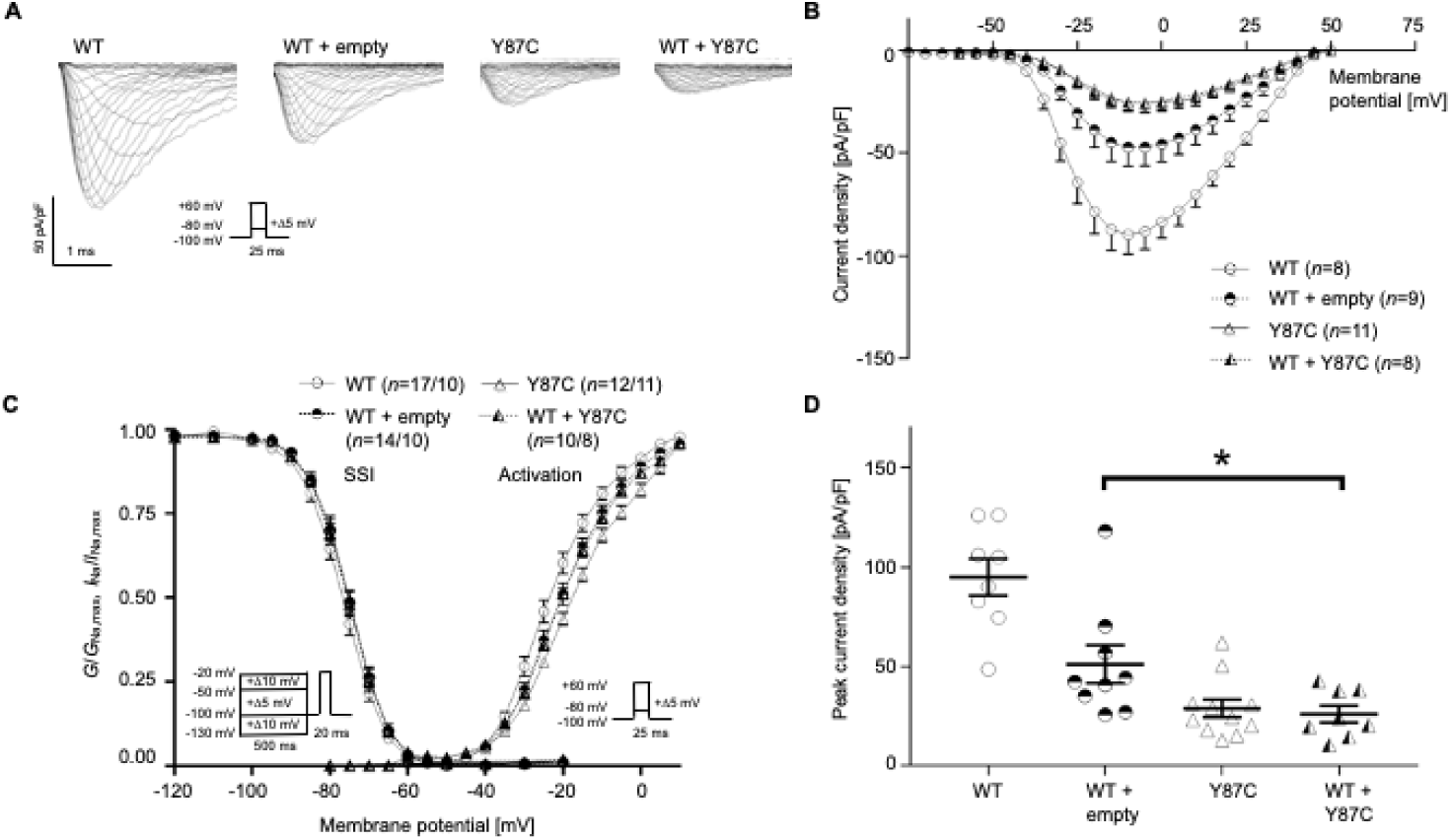
The newly identified BrS-associated Na_v_1.5 variant Y87C exerts dominant-negative effect over WT channels. **(A)** Representative whole-cell *I*_Na_ traces recorded from TsA-201 cells transiently transfected with 100% WT Na_v_1.5, 50% WT + 50% empty vector, 100% Y87C, or 50% WT + 50% Y87C. **(B)** Current density-voltage relationships of the four listed conditions. **(C)** Steady-state inactivation (SSI, left) and activation (right) relationships obtained using the Boltzmann equation. Whole-cell patch clamp protocols are given under the respective curves. **(D)** Peak current densities of each group under unsaturated conditions. Peak current density of WT + Y87C is significantly lower than that of WT + empty vector, indicating that Y87C exerts a dominant-negative effect over WT channels. Values pertaining to the biophysical properties are shown in Table 1. Data are presented as mean ± SEM. *, p < 0.05.

We compared the Y87C variant with the R104 and R121 variants, which were reported to also exert a DNE on WT channel function^2^. Both variants showed no *I*_Na_ in the homozygous state (**Figure 2A-B**), unlike Y87C (**Figure 1A-B**). In the heterozygous state, co-expressing WT + R104W decreased *I*_Na_ peak-current density by ∼55% compared to WT + empty vector, but no significant difference was observed between *I*_Na_ from WT + R121W and WT + empty vector (**Figure 2D**). The V_2/2_ of the activation curves of the different heterozygous conditions did not differ (**Figure 2D, Table 2**).

**Table 2.** Biophysical properties of sodium currents conducted by wild-type, R104W, and R121W Na_v_1.5 channels. Peak sodium current (*I*_Na_) densities are extracted from the respective IV curves. WT, wild type. Data are presented as mean ± SEM and are compared to the respective wild-type values. *, *p* < 0.05; **, *p* < 0.01; **, *p* < 0.005; ***, *p* < 0.001; ****, *p* < 0.0005. Numbers of cells are indicated in parentheses.

| | $I_{Na}$ density | Steady-state inactivation | | Activation | |
| --- | --- | --- | --- | --- | --- |
| | pA/pF | k | $V_{1/2}$ (mV) | k | $V_{1/2}$ (mV) |
| Nav1.5 WT | -86.6±11.4 (n=9) | 5.3±0.2 (n=11) | -73.2±0.5 (n=11) | 6.6±0.2 (n=9) | -22.7±0.6 (n=9) |
| R104W | ND (n=7) | n/a | n/a | n/a | n/a |
| R121W | ND (n=7) | n/a | n/a | n/a | n/a |
| WT + empty v. | -38.4±3.4 (n=5)** | 4.8±0.1 (n=5) | -74.4±0.9 (n=5) | 6.9±0.3 (n=5) | -20.0±0.2 (n=5)** |
| WT + R104W | -23.1±4.6 (n=9)**** | 5.4±0.2 (n=11) | -75.9±0.5 (n=11)** | 7.5±0.3 (n=7)* | -19.5±0.7 (n=7)** |
| WT + R121W | -46.9±3.4 (n=12)** | 5.2±0.2 (n=6) | -72.5±1.3 (n=6) | 7.4±0.3 (n=10)* | -20.3±0.5 (n=10)** |

**Figure 2.**
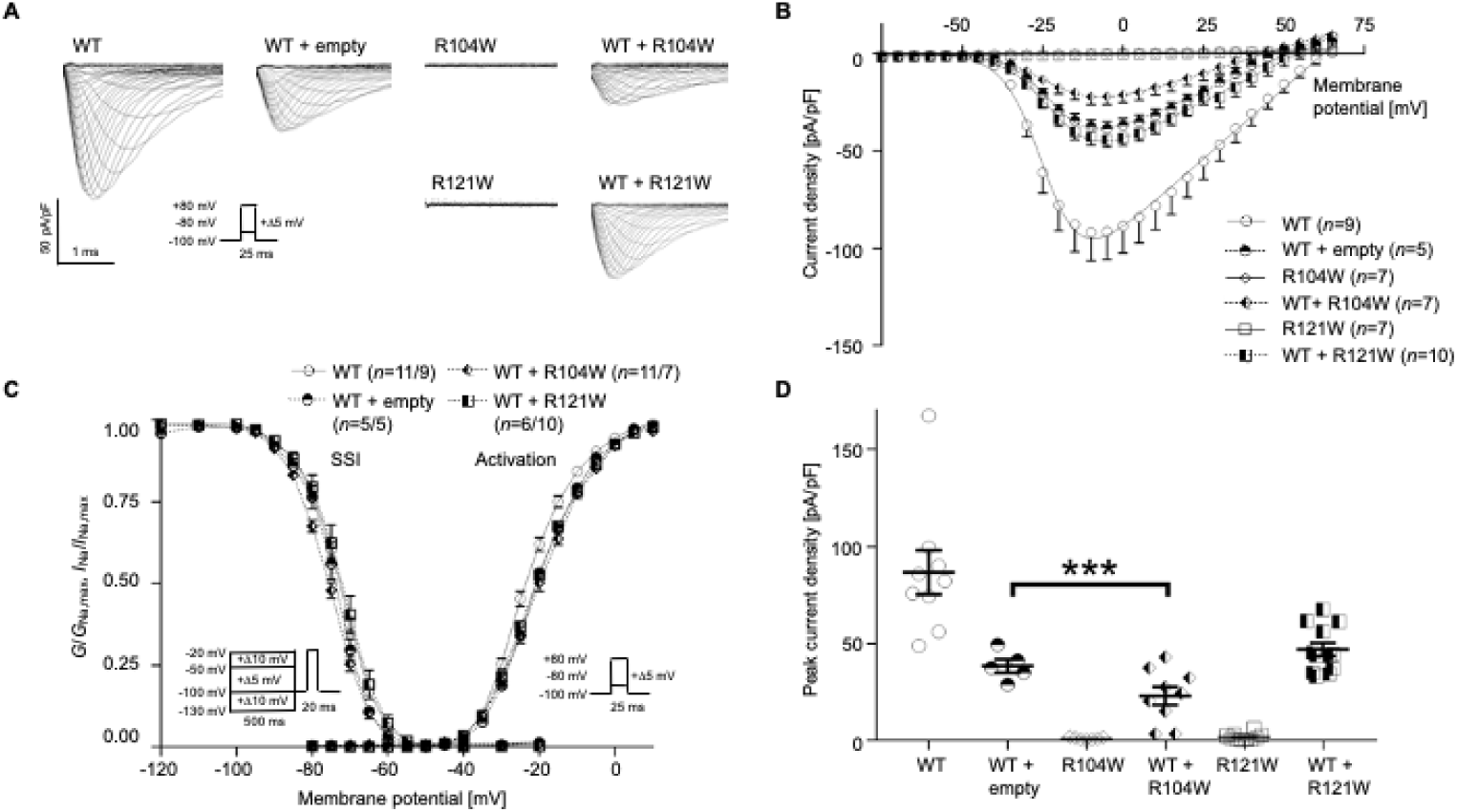
Na_v_1.5 variant R104W but not R121W exerts dominant-negative effect over WT channels. **(A)** Representative whole-cell *I*_Na_ traces recorded from TsA-201 cells transiently transfected with 100% WT Na_v_1.5, 50% WT + 50% empty vector, 100% R104W, 50% WT + 50% R104W, 100% R121W or 50% WT + 50% R121W. **(B)** Current density-voltage relationships of the listed conditions. **(C)** Steady-state inactivation (SSI, left) and activation (right) curves obtained using the Boltzmann equation. Patch clamp protocols are given under the respective curves. **(D)** Peak current densities of each group under unsaturated condition. Peak current density of WT + R104W is significantly lower than that of WT + empty vector, indicating the dominant-negative effect of R104W over WT channels. Values pertaining to the biophysical properties are shown in Table 2. Data are presented as mean ± SEM. ***, *p* < 0.001.

In summary, Na_v_1.5 Y87C and R104W but not R121W negatively regulate WT channel function.

### VARIANT EXPRESSION AND GLYCOSYLATION ARE REDUCED COMPARED TO WT, BUT CHANNEL-ER COLOCALIZATION DOES NOT CHANGE IN VARIANT CONDITIONS

To investigate the mechanisms underlying the reduced or undetectable *I*_Na_ conducted by Y87C, R104W, and R121W compared to WT Na_v_1.5 channels, we investigated protein expression at the cell surface and in whole-cell lysates from TsA-201 cells expressing the aforementioned WT or variant channels. We observed a reduction in Na_v_1.5 variant protein expression compared to WT, both overall and at the cell surface as shown by biotinylation experiments on transfected TsA-201 cells (**Figure 3**). Interestingly, only the high-molecular weight band intensities are reduced both overall and at the surface (**Figure 3B,E**), suggesting that the variant channels are not glycosylated as well as WT channels^2,29^. mRNA expression of variants and WT did not differ in transfected TsA-201 cells (**Supplementary figure 2**).

**Figure 3.**
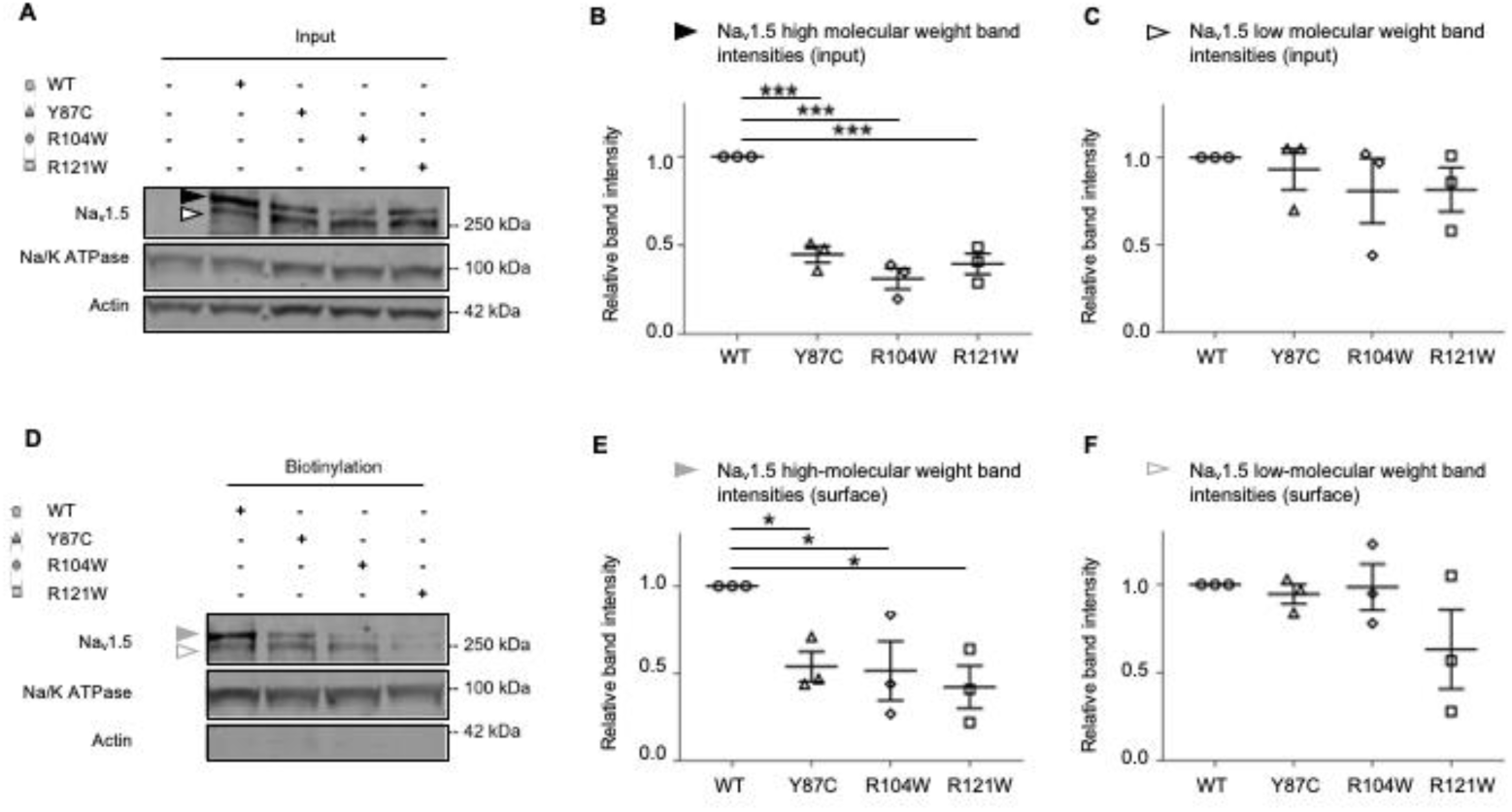
Surface protein expression of all three BrS variants is decreased compared to WT in transfected TsA-201 cells. **(A)** Representative western blot of three independent experiments with whole-cell lysates of TsA-201 cells transiently transfected with WT, Y87C, R104W, or R121W Na_v_1.5 (*n* = 3). Solid and open black arrowheads represent the high- and low-molecular weight bands, respectively. Full blots are shown in Supplementary Figure 4. **(B-C)** Relative protein band intensity of input high- and low-molecular weight bands normalized to Na/K ATPase. Na_v_1.5 WT band intensities are normalized to 1 in each condition. **(D)** Representative western blot of the surface biotinylated fraction of transfected TsA-201 cells (respective whole-cell lysates are shown in **(A)**). Solid and open grey arrowheads represent the high- and low-molecular-weight, respectively. **(E-F)** Relative intensity of surface high- and low-molecular-weight bands normalized to Na/K ATPase. Na_v_1.5 WT band intensities are normalized to 1. Data are presented as mean ± SEM. *, *p* < 0.05; ***, *p* < 0.001.

To assess the intracellular localization of these variants, we transfected rat neonatal cardiomyocytes (RNC) with calreticulin-DsRed ER marker and WT or variant Na_v_1.5 channels and imaged the cells on a confocal microscope with Airyscan (**Figure 4A**). In the homozygous condition, we did not observe any differences in colocalization of WT or variant channels with ER (**Figure 4C**). To mimic the heterozygous condition, we transfected RNC with GFP-WT and FLAG-tagged variant channels, in addition to calreticulin-DsRed (**Figure 4B**). Again, we did not observe any difference in colocalization of WT channels with the ER marker in the presence of WT or variant channels (**Figure 4D**).

**Figure 4.**
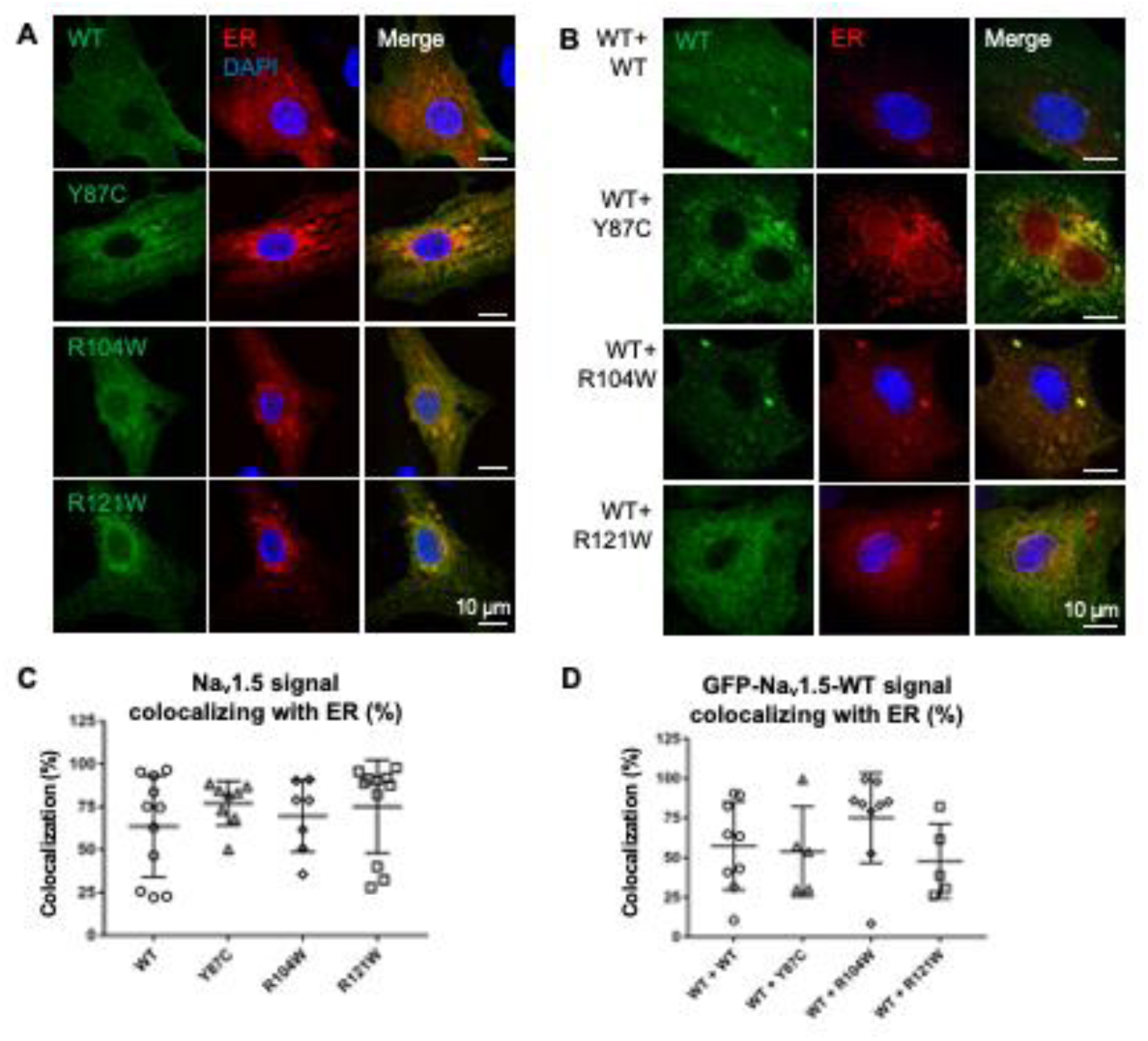
WT and variant channel-ER colocalization is similar in homo- and heterozygous conditions. **(A)** Airyscan microscopy images of RNCs transiently co-transfected with WT or variant flag-Na_v_1.5 (green) and the ER marker calreticulin-DsRed2 (red), representing homologous conditions. Nuclei are stained with DAPI (blue). Right column: merged images. Note that more Na_v_1.5 variant signals colocalize with ER than WT. **(B)** Airyscan microscopy images of RNCs transiently co-transfected with GFP-Na_v_1.5 WT (green), flag-Na_v_1.5 WT or variants (not shown), and calreticulin-DsRed2 (red). Right column: merged images; nuclei are stained with DAPI (blue). **(C)** Quantification of Na_v_1.5-ER colocalization in homozygous conditions. Colocalization is defined as percentage of Na_v_1.5 signal area colocalizing with ER signal area within predefined ROI. **(D)** Quantification of colocalization of WT-Na_v_1.5 with ER in the presence of variants, representing heterozygous conditions. Cell areas in which Na_v_1.5 and ER colocalize are: WT, 57.4 ± 9.3%; Y87C, 54.0 ± 12.6%; R104W, 75.1 ± 9.5%; and R121W, 47.9 ± 10.5%. Scale bar, 10 µm. Data are presented as mean ± SEM.

Taken together, the reduced peak *I*_Na_ density of Y87C, R104W, and R121W compared to WT channels (**Figure 1D, 2D**) correlates with a reduction in surface and overall expression of fully glycosylated variant channels.

### NA_V_1.5 WT NTD ARE WITHIN INTERACTING DISTANCE TO THE NA_V_1.5 WT FULL-LENGTH CHANNEL

To explain how some NTD variants can confer a DNE while others cannot, it is essential to know that the dominant-negative effect is thought to depend on channel dimerization^2^. Based on the observation that co-expressing the NTD with full-length WT Na_v_1.5 increases sodium current^2^, we investigated the possibility that the Na_v_1.5 NTD interacts with full-length channels. We performed proximity ligation assays on COS cells transfected with full-length WT channels and WT NTDs to assess if the Na_v_1.5 NTD is within interacting distance with the whole-length channel *in situ* (**Figure 5A**). **Figure 5B** shows that the WT NTDs are close to full-length WT channels. These results suggest that indeed the Na_v_1.5 WT NTD can interact with full-length WT Na_v_1.5.

**Figure 5.**
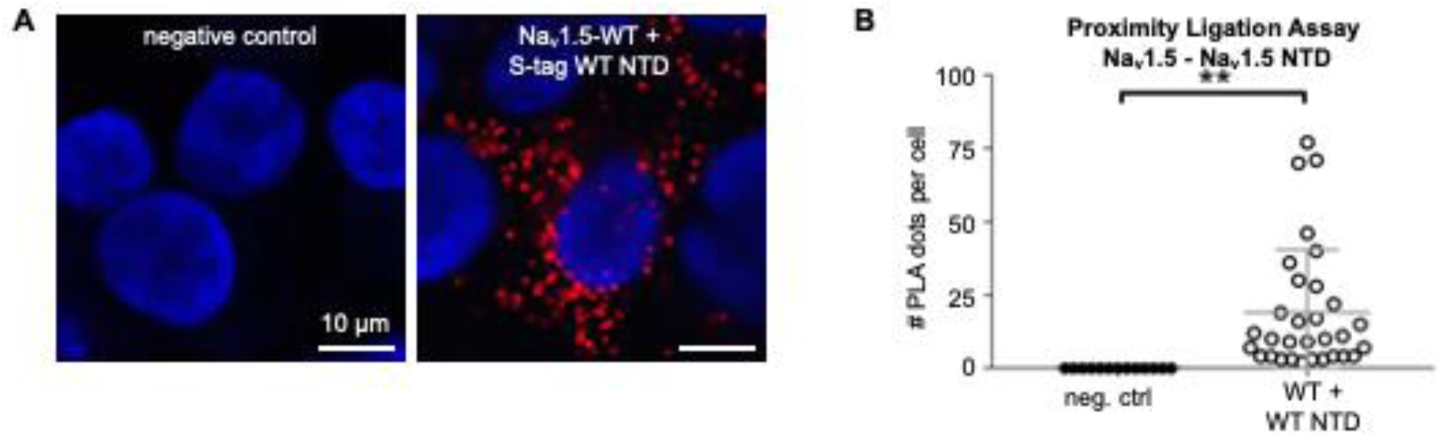
CaM interacts with Na_v_1.5 NTD. **(A)** Representative Duolink^®^ PLA images of COS cells transiently transfected with WT S-tagged Na_v_1.5 NTD. Red dot signals were generated when full-length WT Na_v_1.5 is within 40 nm of WT or variant NTD. Nuclei stained with DAPI in blue. Scale bar, 10 µm. **(B)** Quantitative analysis of the PLA signals. Data are presented as mean ± SEM. **, *p* < 0.005.

### CAM BINDS TO NA_V_1.5 NTD

To explain why the R121W does not exert a DNE while Y87C and R104W do, we hypothesized that the N-terminal domain around position R121 must contain a binding site for a yet-unknown protein. We hypothesized that CaM might be able to bind the Nav1.5 N-terminus based on the following observations.

Firstly, calmodulin binds the NTD of the voltage-gated calcium channel Ca_v_1.2 at two different sites^2,17^, and plays a role in C-terminal dimerization of Ca_v_1.2^2,34^ and Na_v_1.5^2^. Secondly, the NTDs of Na_v_1.5 and Ca_v_1.2 both contain a highly conserved sequence consisting of the residues PIRRA (**Figure 6A,B**), herein the PIRRA motif. In Ca_v_1.2, this motif interacts with calmodulin and calmodulin-binding protein 1 (CaBP1)^2,36^. Moreover, several sequence fragments of the Ca_v_1.2 and Na_v_1.5 NTDs have a high level of sequence identity^2^ (**Figure 6B**).

**Figure 6.**
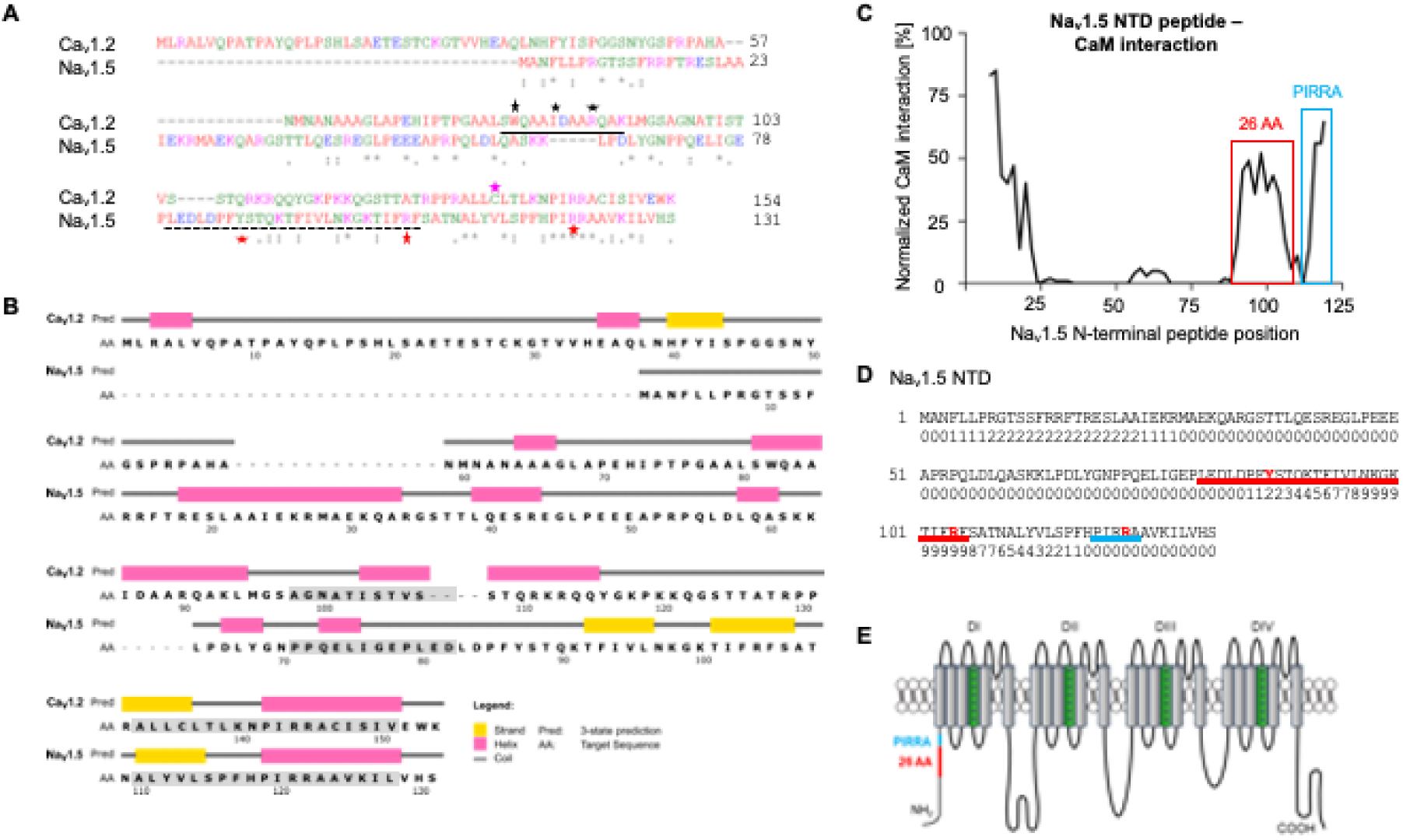
CaM is predicted to bind Na_v_1.5 NTD. **(A)** Sequence alignment of Na_v_1.5 and Ca_v_1.2 NTDs used in this study (https://www.ebi.ac.uk/Tools/msa/clustalo/). The Na_v_1.5 BrS variants studied here are marked with red stars. The 26-AA-long predicted CaM Na_v_1.5 NTD binding site is marked with a dashed black line. Solid black line^17,31^, black stars^32^, and cyan star^16^ indicate CaM binding sites in the Ca_v_1.2 NTD. Residue homologies are scored as follows: asterisk (*), identical; colon (:), strongly similar properties; stop (.), weakly similar properties. Residue color codes: red: small; blue: acidic; magenta: basic; green: hydroxyl, sulfhydryl, amine. **(B)** Secondary structure prediction of Na_v_1.5 and Ca_v_1.2 NTDs (http://bioinf.cs.ucl.ac.uk/psipred/). β-strands (yellow), α-helices (cyan), and coil regions (grey) are indicated. Grey boxes indicate sequences with a high degree of homology (https://blast.ncbi.nlm.nih.gov). **(C)** Normalized interaction between Na_v_1.5 NTD peptides and CaM based on peptide SPOTS arrays. The signal amplitude of the first peptide (AA 1-15) is centered at AA position 8 and the last (AA 112-126) at 121. The interaction was normalized to the strongest signal correspondent to the IQ motif in Na_v_1.5 C-terminal domain. Predicted CaM binding site is marked in red; peptides containing the PIRRA moif in blue. **(D)** Sequence of Na_v_1.5 NTD with predicted CaM binding scores. Scores are normalized on a scale from 0 to 9. A sequence with consecutively high scores suggests the presence of a CaM binding site. The 26 amino acid sequence containing the putative CaM binding site based on peptide SPOTS arrays **(C)** is underlined in red. The PIRRA motif is underlined in blue. The amino acids corresponding to the Y87C, R104W, and R121W variants are given in red. **(E)** Topology of the Na_v_1.5 α-subunit. Red line indicates location of predicted CaM binding sequence, blue line indicates PIRRA motif location. Panel **(E)** adapted from Abriel *et al*.^6^

Firstly, we used the CaM target database to predict the CaM-Nav1.5 NTD binding affinity (Figure 6D).16,32,38 Consecutive high binding scores suggest a CaM binding site, which occurs at amino acid residues ∼94-∼108 (**Figure 6D**), which includes R104, but not Y87 and R121.

We then processed the Na_v_1.5 NTD with a peptide SPOTS array to biochemically assess CaM binding to 15-amino-acid-long N-terminal peptides (**Figure 6C**). We identified a putative CaM-binding sequence comprising 26 amino acids (residues 80-105, **Figure 6C**), which overlapped with the computational prediction (**Figure 6D**) and included Y87 and R104, but not R121.

To validate the putative CaM-binding sequence, we determined whether CaM is close to the Na_v_1.5 NTD *in situ* by performing proximity ligation assays (PLA) on COS cells transfected with S-tagged Na_v_1.5 WT NTD or Δ26 NTD, in which the 26-amino-acid-long putative CaM binding site was deleted (**Figure 7A**). A dot in these images indicates that a Na_v_1.5 NTD and CaM are within 40 nm of each other. We observed that much more CaM proteins are in close proximity to Na_v_1.5 WT NTD than to Δ26 NTD (**Figure 7B**).

**Figure 7.**
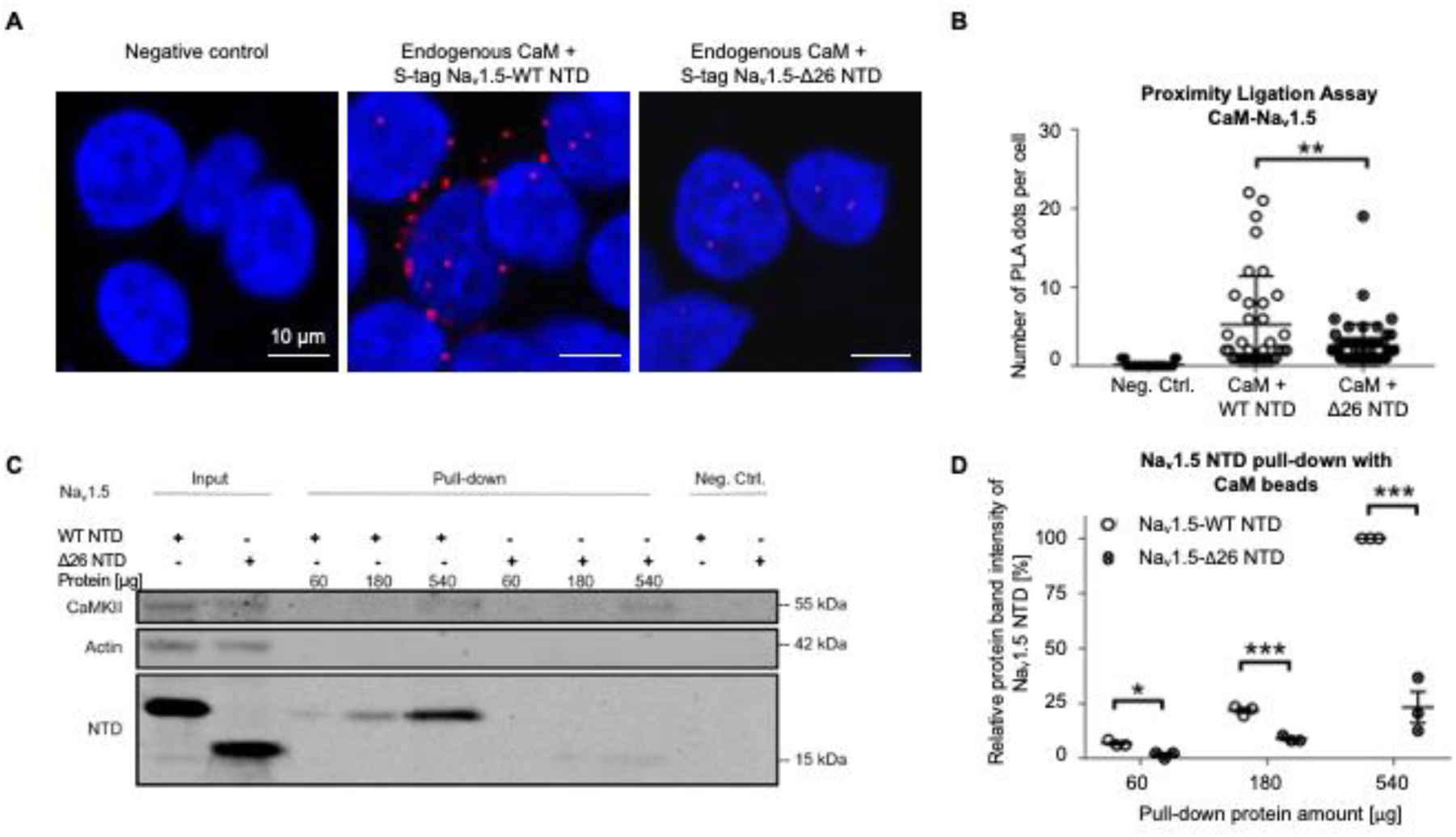
CaM binds to Na_v_1.5 N-terminal domain. (**A)** Representative Duolink^®^ PLA images of COS cells transiently transfected with S-tagged WT NTD, Δ26 Na_v_1.5 NTD, or empty vector as a negative control. Red dots were generated when endogenous CaM and NTD were less than 40 nm apart. Nuclei are stained blue with DAPI. Scale bar, 10 µm. **(B)** Quantitative analysis of the PLA signals. **(C)** Representative western blot of three independent CaM pull-down experiments performed with TsA-201 cells transiently transfected with S-tagged WT or Δ26 Na_v_1.5 NTD. In the latter, the 26 amino acids comprising the predicted CaM-binding sequence (Figure 6) were deleted. We used different amounts of cell lysate (60, 180, 540 µg) to detect CaM-interaction under non-saturating conditions. Full blots are shown in Supplementary Figure 5. **(D)** Relative protein band intensity of pulled-down Na_v_1.5 NTDs. The protein band intensities are normalized to the 540 µg Na_v_1.5 WT NTD condition. Compared to WT, the pulled-down Δ26 NTD relative protein band intensity reduced by ∼5%, ∼12%, or ∼75% with 60, 180, or 540 µg protein lysate, respectively. Data are presented as mean ± SEM. *, *p* < 0.05; **, *p* < 0.005; *** *p* < 0.001.

Next, we performed CaM pull-down assays at the physiological Ca^2+^ concentration of 100 nM to determine if CaM binds to the Na_v_1.5 WT NTD (**Figure 7C**). Lysates from TsA-201 cells transfected with Na_v_1.5 NTD constructs were exposed to CaM-coated or uncoated control sepharose beads. Proteins bound to the beads were eluted and visualized on western blots. The well-established CaM interaction partner CaMKII served as positive control^2^ (**Figure 7C**). We observed that the CaM-coated beads but not the control beads had pulled-down NTD proteins, suggesting that CaM specifically binds to the NTD.

Moreover, deleting the aforementioned 26 amino acids (Δ26) from the NTD greatly reduced CaM binding in all tested conditions with cell lysate amounts containing 60 µg, 180 µg, and 540 µg protein (**Figure 7D**), suggesting that these 26 amino acids comprise a CaM binding site. As less than 540 µg protein did not saturate the western blots (**Figure 7C**), we chose to use 360 µg protein for the following pull-down experiments.

Taken together, these results suggest that CaM interacts with the Na_v_1.5 WT NTD and that this interaction largely depends on 26 residues of the NTD.

### R121W BUT NOT Y87C AND R104W WEAKEN THE NA_V_1.5 NTD-CAM INTERACTION

We next investigated CaM binding to the WT, Y87C, R104W, R121W, and Δ26 Na_v_1.5 NTDs using CaM pull-downs. We observed that the binding of WT, Y87C, and R104W NTDs to the CaM beads was similar, while R121W and Δ26 NTDs did so to a much lesser extent (∼-50%) (**Figure 8A, B**). These results suggest that Δ26 and R121W impair the Na_v_1.5 NTD-CaM interaction, but Y87C and R104W do not. Please note that R121W is outside of the predicted 26 amino acid CaM-binding region (**Figure 3C**), suggesting CaM may bind two NTD sites.

**Figure 8.**
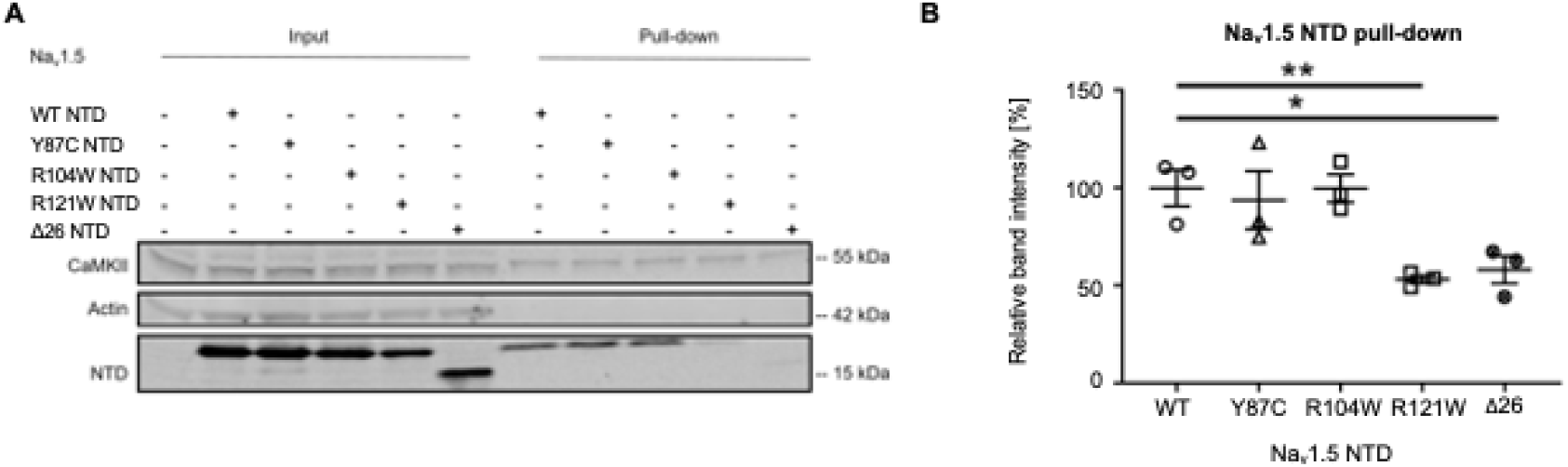
Na_v_1.5 variant R121W weakens the interaction of Na_v_1.5 NTD with CaM. **(A)** Representative western blot of three independent CaM-Na_v_1.5 NTD pull-down experiments using cell lysate equivalent to 360 µg protein per condition from TsA-201 cells transfected with WT, Y87C, R104W, R121W or Δ26 Na_v_1.5 NTD. Full blots including negative controls are shown in Supplementary Figure 6. **(B)** Na_v_1.5 NTD protein band intensities of are normalized to the average WT value. Data are presented as mean ± SEM. ***, *p* < 0.001.

Based on the notion that the homologous Ca_v_1.2 NTD contains a similar arginine to Na_v_1.5-R121, both residing in the PIRRA motif^2,36^, we next determined if mutating the Ca_v_1.2 homologous site R144W also weakens the Ca_v_1.2-CaM interaction. We performed CaM pull-down assays with lysates from TsA-201 cells transfected with Ca_v_1.2-R144W-NTD or Ca_v_1.2-WT-NTD (**Supplementary figure 3A, B**). We found that CaM binds to Ca_v_1.2 WT NTD, consistent with previous reports^2,17^, while R144W partly abolished the interaction with CaM by 66.0 ± 7.7% (**Supplementary figure 3A, B**), which is expected as R144 lies within one of the two N-terminal sequences previously associated with CaM binding^2^.

In summary, in the Na_v_1.5 NTD, both the predicted 26 amino acid sequence and R121 are involved in CaM binding. Similarly, the Ca_v_1.2 NTD R144 site is homologous to R121 and required for CaM binding.

## DISCUSSION

In this work, we demonstrated that CaM binds the N-terminus of Na_v_1.5, which resembles CaM binding to the NTD of Ca_v_1.2^2,17,33,34^. This interaction seems to be crucial for the dominant-negative effect, in which a Na_v_1.5 NTD variant identified in BrS patients negatively regulates wild-type channel function. Specifically, we identified in probands of a Russian family the new natural Brugada syndrome variant Y87C in the Na_v_1.5 NTD and showed that Y87C-Na_v_1.5 channels exert a DNE on WT channels when co-expressed in TsA-201 cells.

We also showed that natural BrS variants in the Na_v_1.5 NTD do not consistently show a DNE: Y87C and R104W do, while R121W does not. To explain this discrepancy, we showed that CaM binding to Na_v_1.5 NTD partly depends on amino acids 80-105 (a 26-AA-long sequence) or residue R121. Thus, the DNE of the tested NTD BrS variant correlates with CaM-NTD interaction strength. Moreover, all tested BrS variant full-length channels show reduced expression and glycosylation at the cell surface than wild-type channels. We schematically summarize the functional consequences of the BrS variants in **Figure 9**.

**Figure 9.**
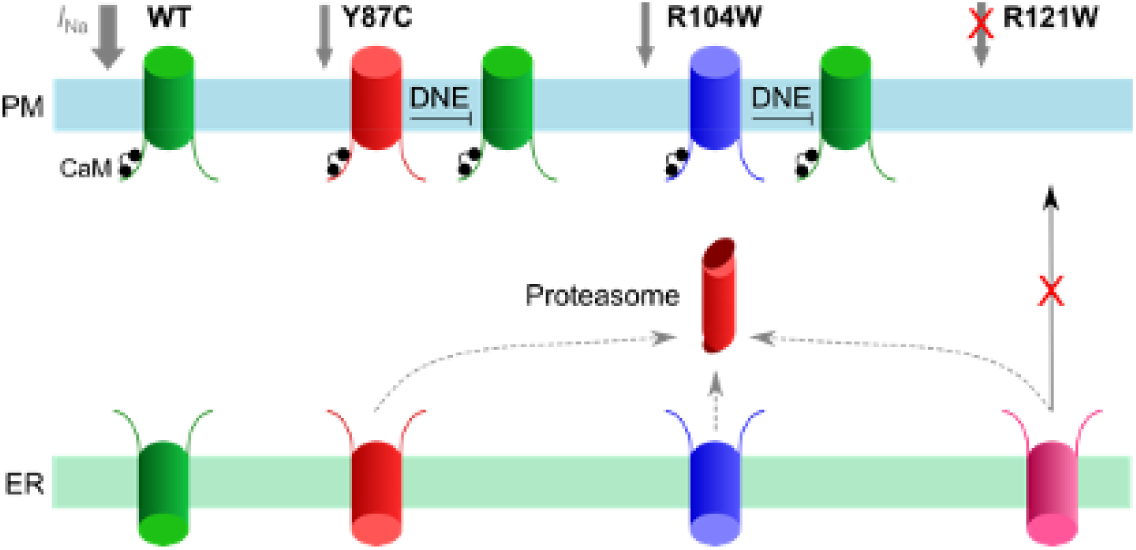
Working model of the CaM-Na_v_1.5 NTD interactions and Na_v_1.5 variants in homozygous conditions. Na_v_1.5 WT channels (green) are readily trafficked from the ER to the plasma membrane and conduct a normal *I*_Na_ density (grey). Y87C (red) or R104W (blue) channel proteasomal degradation (red) is increased and *I*_Na_ is decreased compared to WT, while exerting a DNE on variant channels. R121W channels (pink) do not conduct a detectable sodium current and do not exert a DNE on wild-type channels. CaM (black) interacts with WT, Y87C, and R104W channels but not with R121W.

### CAM BINDS TO THE NA_V_1.5 N-TERMINAL DOMAIN

We established the CaM-Na_v_1.5 NTD interaction based on *in silico* modeling, SPOTS assay, pull-down assays and *in situ* hybridization assays using heterologous expression systems transiently expressing Na_v_1.5 NTD constructs. This interaction partly depends on R121 and amino acids 80-105. We moreover showed that full-length Na_v_1.5-Na_v_1.5 NTD interaction is reduced when amino acids 80-105 are deleted or R121 is mutated (**Figure 8**). Whether CaM interacts with the Na_v_1.5 NTD *in vivo*, and whether this interaction is involved in Na_v_1.5 dimerization, remain open questions. We show in COS cells that the Na_v_1.5 NTD is within interacting distance (<40 nm) of full-length Na_v_1.5 (**Figure 5**); thus, we may hypothesize that Na_v_1.5 channels interact *in vivo* in a CaM-mediated manner. We showed that the natural BrS variant R121W in the Na_v_1.5 homologous PIRRA motif diminishes interaction of NTD with full-length Na_v_1.5. The same was observed when deleting amino acids 80-105, which suggests that CaM may have two binding sites, potentially one for each lobe. At the Na_v_1.5 DII-III linker, distinct binding sites have already been described for the CaM C- and N-lobe^2,41^. Future research efforts should be directed towards specifying the intricacies of the CaM-Na_v_1.5 NTD binding. Besides potentially playing a role in channel dimerization, CaM binding to the Na_v_1.5 NTD may have other effects. At the Na_v_1.5 C-terminal IQ motif, for instance, CaM interaction reduces the late sodium current^2^, and affects voltage-dependence of activation in an calcium-dependent manner^2,43,44^. Given the multiple CaM binding sites at a full-length Na_v_1.5 channel, the stoichiometry of Na_v_1.5:CaM remains to be determined. Previous FRET experiments have shown a 1:1 stoichiometry^2^; however, the Na_v_1.5 N-terminus was not included in the respective Na_v_1.5 peptides.

Besides CaM, other Na_v_1.5 NTD binding partners may play a role in Na_v_1.5 dimerization and the DNE. α1-syntrophin has been shown to bind the three C-terminal residues of Na_v_1.5 and serine-20 of the Na_v_1.5 NTD^2,46^, where it plays a role in the chaperone effect of the NTD, in which co-expressing the Na_v_1.5 NTD with full-length Na_v_1.5 (or with Kir2.1 and −2) leads to higher *I*_Na_ (or *I*_K1_) compared to full-length channel expression alone in CHO cells^2^. Together with the notion that α1-syntrophin is a critical mediator for Na_v_1.5 anchoring to the cytoskeleton^2-49^, these findings suggest that α1-syntrophin may too mediate channel-channel interactions and/or clustering. However, the α1-syntrophin-binding residue serine-20 is not affected by any of the NTD constructs used in this study.

### THE DOMINANT-NEGATIVE EFFECT OF NA_V_1.5 VARIANTS

Contrary to a previous report^2^, we show that R121W full-length channels do not exert a DNE on wild-type channels, while R104W and the newly identified variant Y87C do (**Figure 1, 2**). We also could not reproduce the positive shift in activation of the sodium current when R104W and R121W channels are co-expressed with WT channels as Clatot *et al.* observed^2^. These discrepancies may lie in experimental conditions (TsA-201 cells and untagged *SCN5A* constructs in our study versus HEK293 cells and GFP-tagged *SCN5A* constructs used by Clatot *et al.*^2^). Experiments in hiPSCs from variant-carrying BrS patients or in mouse models heterozygous for R104W, R121W, or Y87C could shed light on this discrepancy in a more native or *in vivo* system.

Although our data do not give direct evidence for Na_v_1.5-Na_v_1.5 dimerization, we may hypothesize that the observed DNE of R104W and Y87C BrS variants depends on Na_v_1.5 NTD dimerization based on the following observations. Firstly, partly abolishing the CaM interaction by mutating R121 correlated with the absence of the DNE (**Figure 2**). Secondly, CaM plays a role in Na_v_1.5 and Ca_v_1.2 dimerization at their C-termini^2,33-36^. This may be extrapolated to the Na_v_1.5 NTD but remains to be investigated. Thirdly, Na_v_1.5 dimerization mediated by 14-3-3 at the DI-II linker has been shown to be crucial for Na_v_1.5 DNE as inhibiting 14-3-3 binding abolished the DNE^2^. In addition to 14-3-3, Mercier *et al.* reported that the sodium channel β^2^-subunit was required for the DNE^2^. Indeed, the DNE of Na_v_1.5 loss-of-function variants may involve many more molecular determinants. Partly underlying this gap in knowledge is the notion that research groups functionally characterizing Na_v_1.5 variants rarely co-express WT with variant channels; therefore, literature on the Na_v_1.5 DNE is relatively scarce^2^. How CaM, 14-3-3, the β^2^-subunit, and potential other Na_v_1.5 interacting proteins interdependently or independently establish the DNE, and which known Na_v_1.5 variants do and do not exert a DNE, are exciting venues for future research.

The hypothesis that channel di- or multimerization underlies the DNE of variant Na_v_1.5 is further indirectly supported by the multimerization of other voltage-gated channels. Ca_v_1.2 channels in the heart for instance have been shown to be functionally and physically coupled in a CaM- and Ca^2+^-mediated manner^2^. This coupling is increased by ß-adrenergic stimulation^2^. In the brain, functional coupling has been shown for BK and Ca_v_1.3 channels, and cooperative gating has been suggested to play a role in short-term memory^2,53^. Lastly, voltage-gated proton channels that are expressed in many organisms and tissues are also shown to gate cooperatively^2-56^. The body of knowledge regarding channel-channel interactions however is still in its infancy, and the roles of these interactions in the pathogenesis of cardiac arrhythmias remain to be uncovered.

### DECREASED NA_V_1.5 PROTEIN EXPRESSION AND ITS SUBCELLULAR LOCALIZATION

We observed a reduction in protein expression of the fully glycosylated form of Y87C, R104W, or R121W channels compared to wild-type, both at the surface and in whole-cell lysates, whereas expression of the core-glycosylated form did not change (**Figure 3**). Functionally, a reduction in fully glycosylated Na_v_1.5 channels has been shown to change voltage-dependency of Na_v_1.5^2^; however, we have not observed this effect in our own functional experiments (**Figure 1,2**). We investigated whether this could be explained by variants retaining wild-type channels in the ER, but in the presence of variant channels, wild-type channels showed similar degrees of co-localization with the ER (**Figure 4**). mRNA expression was the same between groups (**Supplementary figure 2**), indicating that variability in transfection rate, gene transcription, or mRNA processing do not underlie the differences in protein expression. Rather, we expect that the reduction in fully glycosylated variant Na_v_1.5 may be the result of increased proteasomal degradation. Mercier *et al.* have shown that the fully glycosylated band represents Na_v_1.5 channels that have followed the pathway from ER to Golgi and the plasma membrane, whereas core-glycosylated channels are transported from the ER straight to the plasma membrane^2^. Thus, we hypothesize that fully glycosylated BrS variant channels having passed through the Golgi seem more susceptible to proteasome-mediated degradation. The underlying mechanisms however remain unclear and offer exciting venues for future study.

## CONCLUSION

Based on our experimental data, we can conclude that the novel naturally occurring Y87C variant is likely directly linked to the BrS phenotype of the probands. Moreover, we show novel calmodulin binding sites at the Na_v_1.5 N-terminal domain, in conjunction with its putative role in the dominant-negative effect of natural Brugada syndrome variants. These results need to be validated *in vivo* and the intricacies of CaM-Na_v_1.5 NTD binding remain to be unraveled.

## ACKNOWLEDGMENTS

The authors sincerely thank Regula Flückiger-Labrada for her excellent work in isolating rat neonatal cardiomyocytes, Anne-Flore Hämmerli for technical assistance, Dr. Nathalie Neyroud and Dr. Jin Li for fruitful discussions, and Dr. Kali Tal for her editing of a previous version of the manuscript. The authors also acknowledge the contribution of the Microscopy Imaging Center (MIC), University of Bern.

## AUTHOR CONTRIBUTIONS

Z.W. and H.A. designed this research project. Z.W., A.H., V.S., A.S., and D.R.K performed the experiments. E.V.Z. provided the ECG and clinical description. Z.W., S.V., A.H., and D.R.K. analyzed the data. Z.W. and S.V. made figures and tables. Z.W. and S.V. drafted and edited the manuscript. Z.W., S.V., D.R.K., G.P., and H.A. critically reviewed the manuscript.

## ABBREVIATIONS

BrS: Brugada syndrome
CaM: Calmodulin
Ca_v_ channels: Voltage-gated calcium channels
CTD: C-terminal domain
DNE: Dominant negative effect
ECG: Electrocardiogram
ICD: Implantable cardioverter defibrillator
I_Na_: Sodium current
LQT: Long-QT syndrome
MI: Myocardial infarction
Na_v_ channels: Voltage-gated sodium channels
NTD: N-terminal domain
PLA: Proximity Ligation Assay
RNC: Rat neonatal cardiomyocytes

## SUPPLEMENTARY FIGURES

**Supplementary Figure 1.**
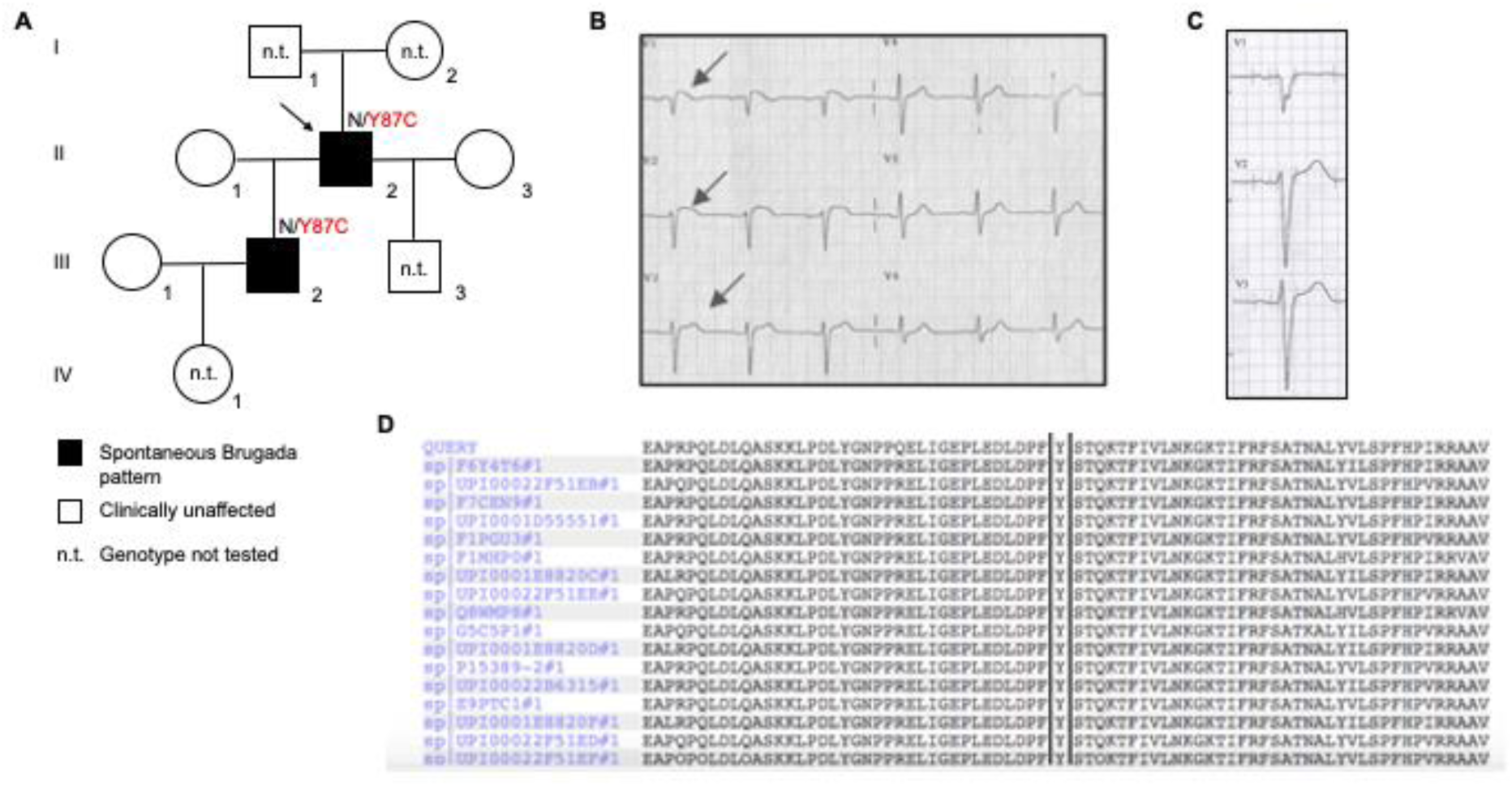
Prevalence of Brugada Syndrome and the Y87C Nav1.5 variant in the Russian proband family. **(A)** Pedigree of the proband family. The proband is marked with a black arrow. Black squares indicate family members with BrS; empty symbols indicate clinically unaffected family members. n.t: genotype not tested. **(B)** ECG from proband II.2 recorded at 49 years old shows a spontaneous Brugada pattern. Heart rate was 75 bpm, QTc-interval 383 ms, and PR-interval 240 ms. An ST-elevation > 2mm is observed in V1-V3 with a negative T-wave in V1. (C) ECG from III.3 recorded at 13 years old shows no Brugada pattern. ECG registered at 50 mm/s, amplitude 10 mm/mV. (D) Multiple sequence alignment showing the highly conserved p.87Y position in Nav1.5 across different species.

**Supplementary Figure 2.**
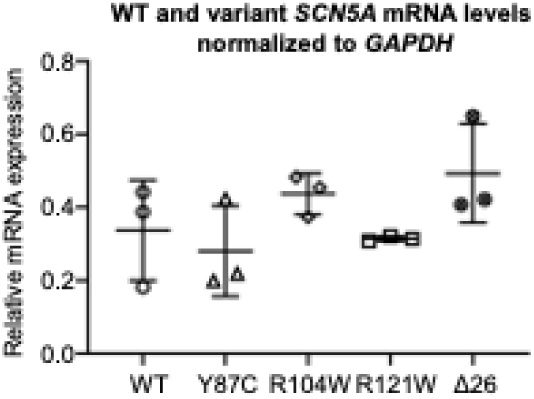
Na_v_1.5 WT- and variant-encoding mRNA expression levels were similar. **(A)** RT-qPCR data of *SCN5A* mRNA expression in TsA-201 cells transiently transfected with WT-, Y87C-, R104W-, and R121W-Na_v_1.5-encoding cDNA normalized to *GAPDH* showed no difference between the conditions.

**Supplementary Figure 3.**
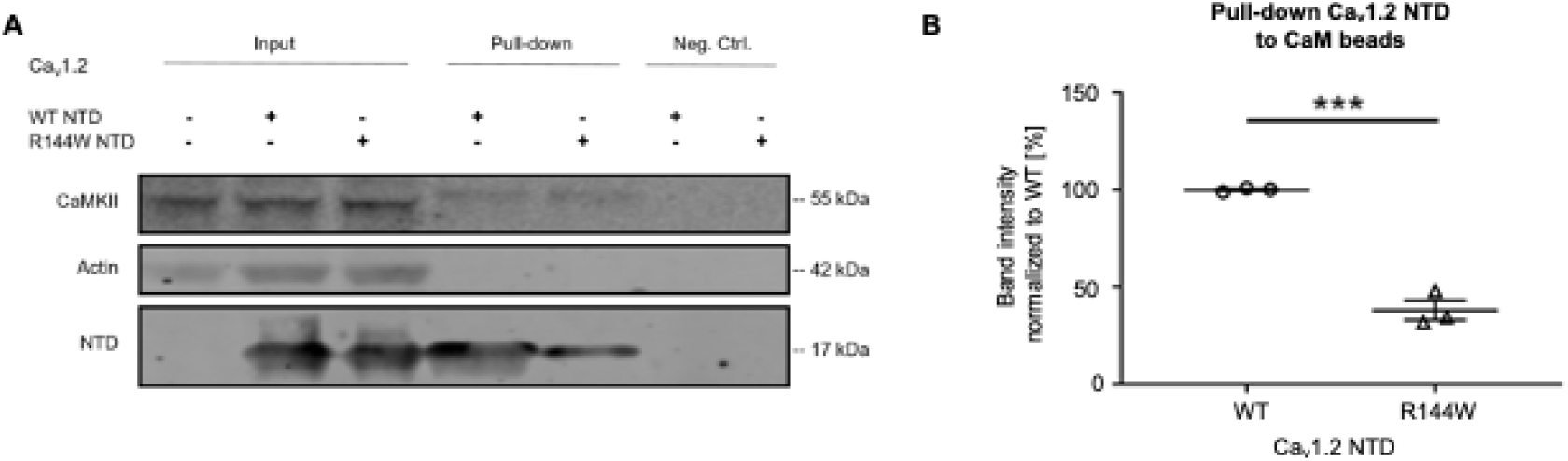
The R144W mutation in Ca_v_1.2 weakens the Ca_v_1.2 NTD-CaM interaction. **(A)** Representative western blot of three independent CaM-Ca_v_1.2 NTD pull-down experiments performed with 360 µg TsA-201 cell lysate. Full blots are shown in Supplementary Figure 7. **(B)** Relative protein band intensity of Ca_v_1.2 NTD normalized to endogenous CaMKII. The Ca_v_1.2-WT NTD band intensities are normalized to 1. Data are presented as mean ± SEM. ***, *p* < 0.001.

**Supplementary Figure 4.**
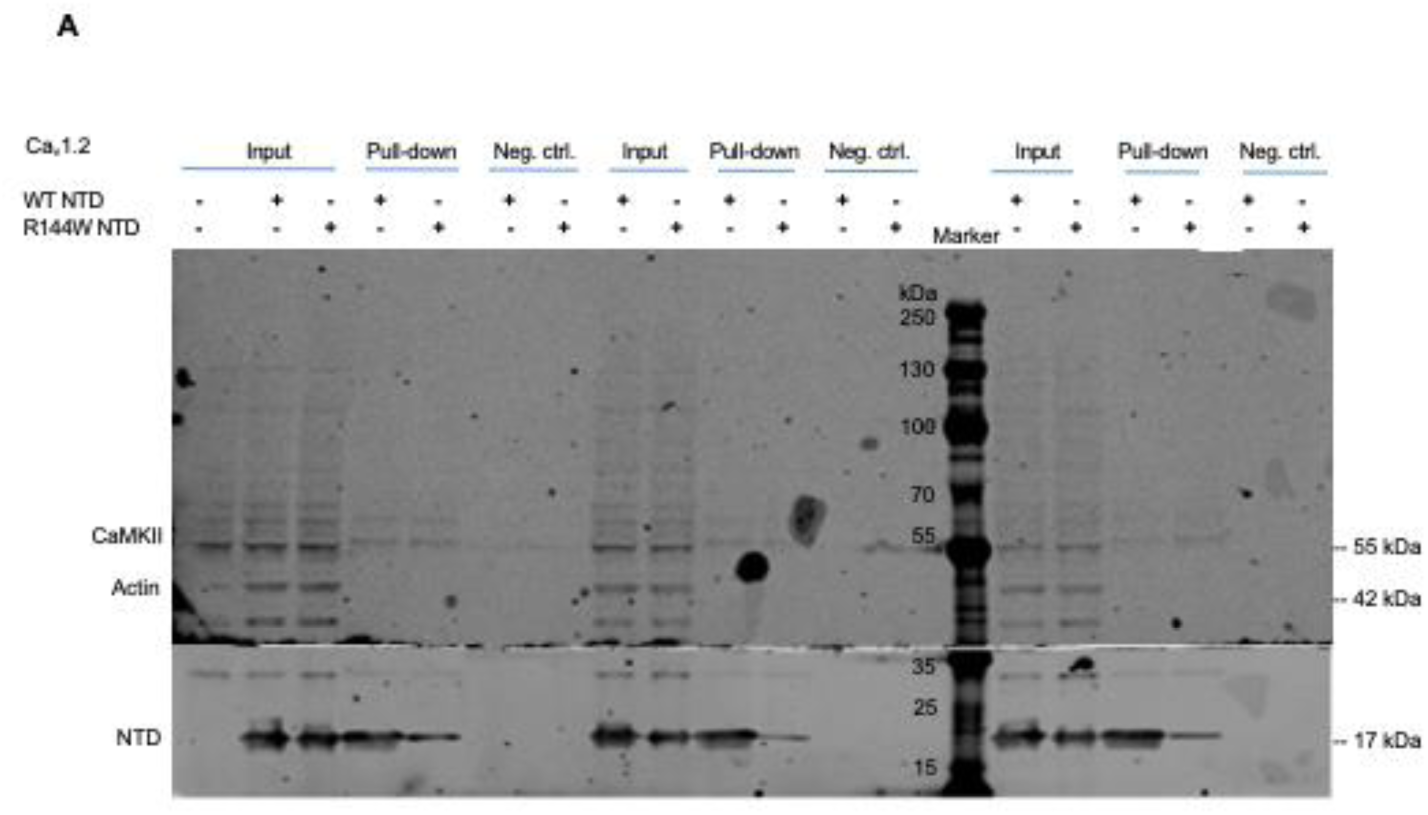
Full western blots of three independent biotinylation experiments in TsA-201 cells transiently transfected with Na_v_1.5 WT, Y87C, R104W, and R121W.

**Supplementary Figure 5.**
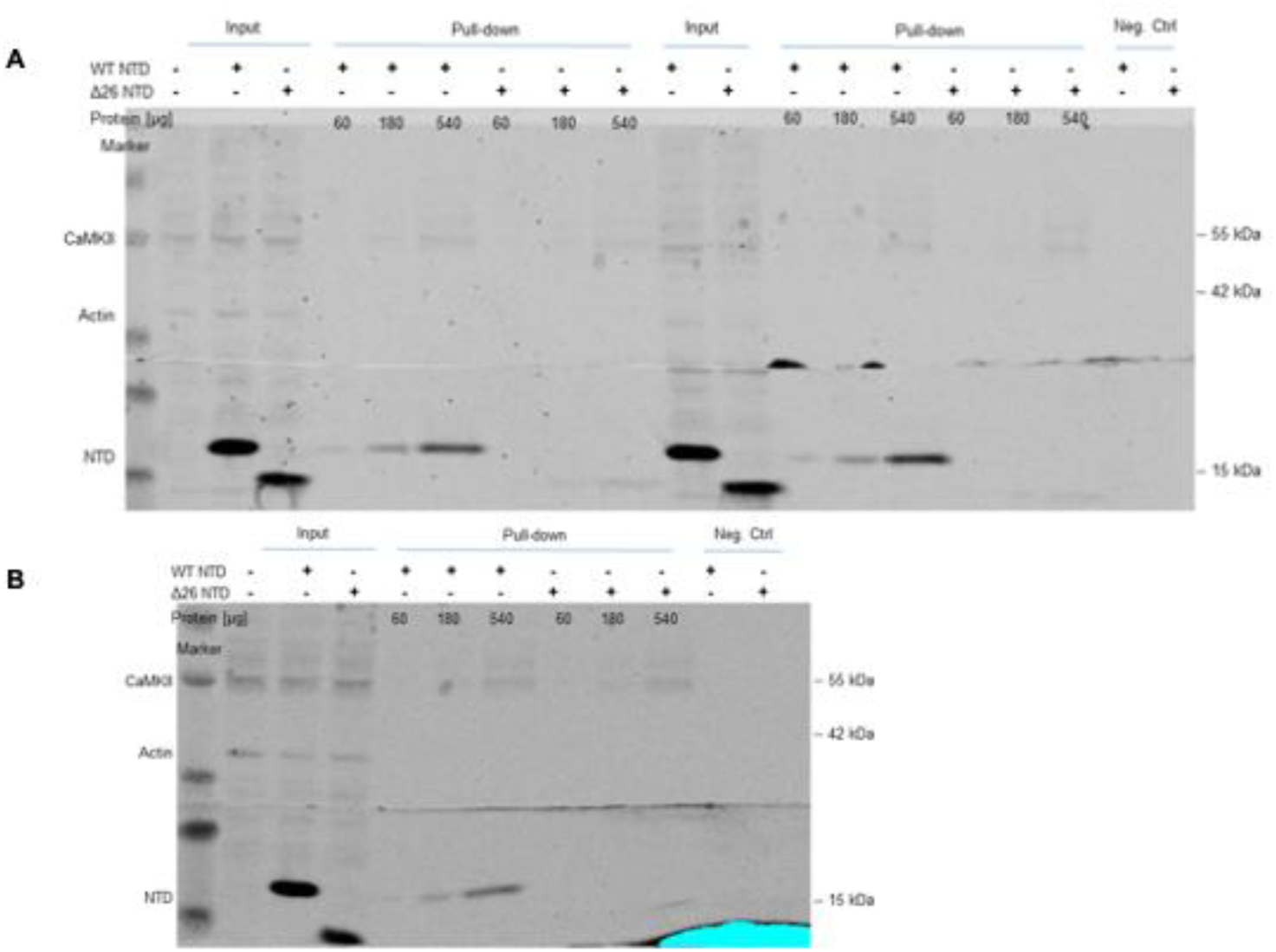
Full western blots of three independent CaM pull-down experiments in TsA-201 cells transiently transfected with S-tagged Na_v_1.5 WT and Δ26 NTD using various amounts of protein.

**Supplementary Figure 6.**
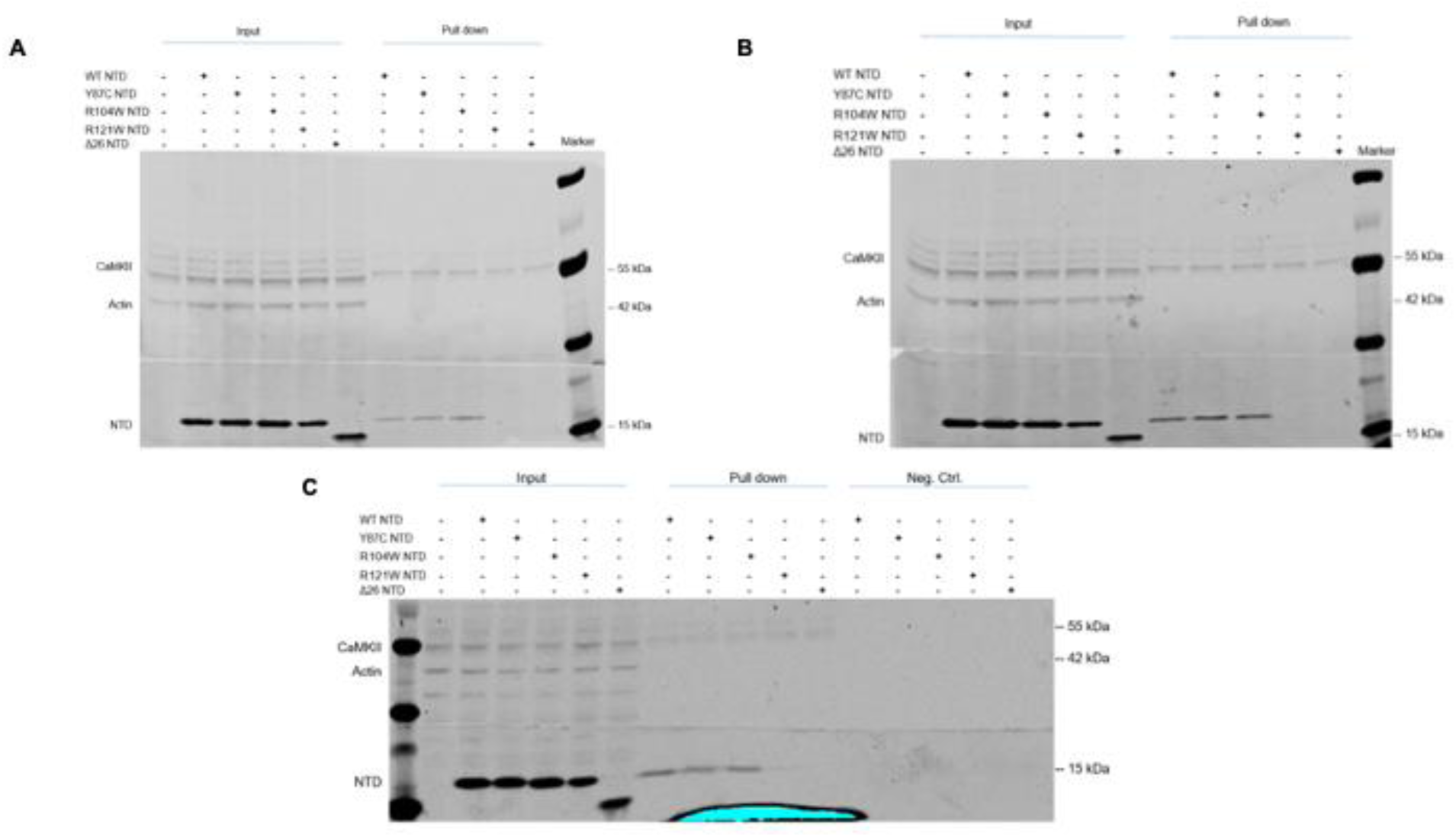
Full western blots of three independent CaM pull-down experiments with lysates equivalent to 360 µg per condition from TsA-201 cells transiently transfected with S-tagged Na_v_1.5 WT, Y87C, R104W, R121W, and Δ26 NTD.

**Supplementary Figure 7.**
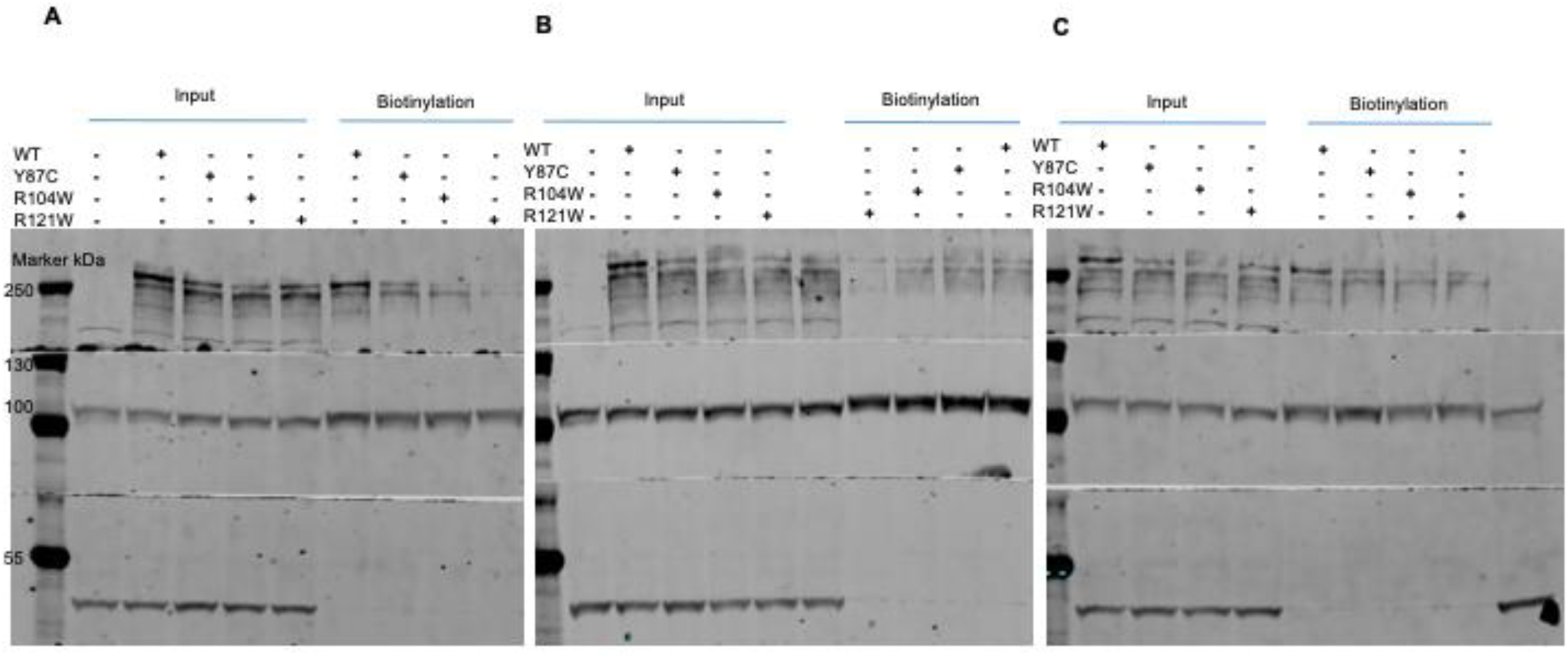
Full western blots of three independent CaM pull-down experiments with lysates equivalent to 360 µg per condition from TsA-201 cells transiently transfected with S-tagged Ca_v_1.2 WT and R144W NTD.

